# Integrated analysis of directly captured microRNA targets reveals the impact of microRNAs on mammalian transcriptome

**DOI:** 10.1101/672469

**Authors:** Glen A. Bjerke, Rui Yi

## Abstract

MicroRNA (miRNA)-mediated regulation is widespread, relatively mild but functionally important. Despite extensive efforts to identify miRNA targets, it remains unclear how miRNAs bind to mRNA targets globally and how changes in miRNA levels affects the transcriptome. Here we apply an optimized method for simultaneously capturing miRNA and targeted RNA sites to wildtype, miRNA knockout and induced epithelial cells. We find that abundantly expressed miRNAs can bind to thousands of different transcripts and many different miRNAs can regulate the same gene. Although mRNA sites that are bound by miRNAs and also contain matches to seed sequences confer the strongest regulation, ∼50% of miRNAs bind to RNA regions without seed matches. In general, these bindings have little impact on mRNA levels and reflect a scanning activity of miRNAs. In addition, different miRNAs have different preferences to seed matches and 3’end base-pairing. For a single miRNA, the effectiveness of mRNA regulation is highly correlated with the number of captured miRNA:RNA fragments. Notably, elevated miRNA expression effectively represses existing targets with little impact on newly recognized targets. Global analysis of directly captured mRNA targets reveals pathways that are involved in cancer, cell adhesion and signaling pathways are highly regulated by many different miRNAs in epithelial cells. Comparison between experimentally captured and TargetScan predicted targets indicates that our approach is more effective to identify *bona fide* targets by reducing false positive and negative predictions. This study reveals the global binding landscape and impact of miRNAs on mammalian transcriptome.

## Background

MicroRNAs (miRNAs) are small non-coding RNAs that repress gene expression through sequence-specific mRNA target binding, resulting in mRNA degradation or repression of translation [1]. The majority of mammalian genes are thought to be regulated by miRNAs, suggesting that miRNA-mediated gene expression regulation is widespread and functionally important [2,3]. However, understanding the function of an individual miRNA or miRNA family has been hampered by the fact that a single miRNA may regulate many genes and a single gene can be regulated by many different miRNAs [4]. In addition, a single miRNA:RNA pairing usually results in mild regulation of gene expression, indicating that multiple interactions are necessary to impart strong repression, either against one gene or multiple components within a pathway [5]. This also indicates that to comprehensively understand the effect of a miRNA or miRNA family, the regulation of the entire transcriptome has to be quantitatively measured as opposed to one or a few targets of interest in a cell type-specific manner [6,7]. Furthermore, when the expression levels of miRNAs elevate in conditions such as normal development, stem cell self-renewal and differentiation, pathogenesis or through experimental manipulation, it is unclear whether the increased miRNA expression causes stronger binding and repression to canonical targets or if this results in *de novo* miRNA:RNA interactions with new targets.

miRNA:RNA interaction is mediated by partial base pairing between miRNA and RNA sequences. During miRNA biogenesis in mammals, a single-stranded mature miRNA is loaded onto one of the four Argonaute (Ago) proteins, usually determined by the protein abundance [8], to form the RNA induced silencing complex (RISC) [9]. Structural, biochemical and computational studies have all demonstrated that the sequence at the 5’ end of miRNAs, often termed the seed, are most critical to miRNA:RNA interactions [10] although miRNA:RNA interactions via the 3’ regions of miRNAs also likely play a role [11]. Over the past fifteen years, extensive efforts have been dedicated to develop computational tools, which are generally based on identifying mRNA regions with a seed match, to identify miRNA targets. Prominent algorithms such as TargetScan have become a major resource for miRNA target prediction [2]. However, computational predictions generally cannot count for variable cellular contexts or detect non-perfectly matched targets and they also lack the ability to distinguish functionally important target genes or pathways, usually due to a large number of predicted targets [12]. Given the difficulty of predicting targets, many experimental methods for determining targets have been developed. For example, HITS-CLIP method has been established to identify miRNA target sites that are crosslinked by UV radiation and associated with Ago proteins [13]. Because miRNA recognized target sites are bound and protected by Ago proteins, they are subsequently recovered from Ago immunoprecipitation and deeply sequenced for identification of miRNA targets. PAR-CLIP was further developed to improve the crosslink efficiency of HITS-CLIP by incorporating 4-thiouridine into the RNA in cultured cells [14]. Both methods, however, do not identify miRNAs that mediate the recognition of mRNA target sites and often detect large mRNA regions that are associated with Ago proteins. As a result, extensive bioinformatic analysis, often relying on the presence of miRNA seed matches (6mer to 8mer sequences) within mRNA sequences, has to be used to identify miRNA binding sites and assign them to specific miRNAs (Chi et al. 2009 and Hafner et al. 2010). Because of the prevalence of 6mer to 8mer sequences in mammalian transcriptome, such an identification often generates false positives and also cannot assign mRNA regions without any match to canonical miRNA seed regions. To overcome these difficulties, the CLASH technique was reported to identify miRNA targets by sequencing ligated miRNA:mRNA chimeras, which allows the identification of single miRNA and its associated mRNA site regardless of the presence of seed matches [15,16]. The low frequency of miRNA:mRNA chimeric reads in the original CLASH study was then improved upon by the CLEAR-CLIP method that adds an additional ligation step to enhance miRNA:mRNA ligation [11]. This technique allows identification of *bona fide* miRNA targets through direct capture and sequencing of miRNA:RNA pairs that were ligated while still in the RISC and removes the imperfect bioinformatic prediction that assigns a miRNA to an mRNA site as required by HITS-CLIP [11]. However, it remains unclear whether CLEAR-CLIP can distinguish the strong and weak miRNA target sites and how CLEAR-CLIP identified miRNA:mRNA interactions reflect miRNA-mediated mRNA regulation.

In this study, we utilize improved experimental conditions that allow more efficient and unbiased RNA ligation [17] to enhance the ability of CLEAR-CLIP to capture miRNA:RNA pairs. By using randomized adapters during the ligation steps, we further improve the quantitative performance of CLEAR-CLIP. To demonstrate the effectiveness of the optimized CLEAR-CLIP, we apply this method to study the miR-200 family. The five members of miR-200s differ by a single nucleotide within the seed region and also have variable sequences at the 3’end (Fig. S1a). Thus, a highly accurate target identification method is required to distinguish how each member of the family recognizes and regulates their targets. In addition to wildtype cells, we use primary epithelial cells isolated from miR-200 double knockout mice, in which all five members of the miR-200 family are deleted, and miR-200 inducible mice, in which three members of the family are induced [18]. These data are also combined with RNA-seq of miR-200 induced cells, allowing determination of which miRNA:RNA interactions are functional. We further validate our method with miR-205, one of the most highly expressed miRNAs in epithelial stem cells [19]. The lessons learned from these individual miRNAs are then applied to the miRNA pathway globally, generating new knowledge of miRNA regulated networks and how they control pathways involved in cancer, cell adhesion and signaling in epithelial cells of the skin. Together, this work has established an experimental method to accurately capture all miRNA targets in a miRNA- and target site-specific manner.

## Results

### CLEAR-CLIP identifies miRNA site-specific interactions on a genomic scale

We previously generated mouse skin specific models for double knockout of the two miR-200 clusters (*Krt14-Cre/miR-200c:141^fl/fl^/miR-200b:200a:429^−/−^*, hereafter referred to as DKO) and transgenic miR-200 inducible overexpression (*Krt14-rtTA/pTRE2-miR-200b:200a:429*, hereafter referred to as miR-200 Tg)[18]. We performed CLEAR-CLIP on mouse keratinocytes from nine control samples, six miR-200 DKO samples and three miR-200 Tg samples, allowing us to validate loss or gain of CLEAR-CLIP signals (samples detailed in Table S1). We also combined the CLEAR-CLIP results with RNA-Seq data from miR-200 Tg and miR-205 induced mouse keratinocytes (*Krt14-rtTA/pTRE2-miR-205*, hereafter referred to as miR-205 Tg) to more thoroughly characterize mRNA targeting and expression by the miR-200 family, miR-205 and the entire miRNA pathway. The miR-200 Tg keratinocytes used for RNA-Seq were induced with doxycycline for 24 hours, resulting in overexpression of the miR-200b/a/429 cluster approximately 15-fold (Fig. S1b). miRNAs can affect both mRNA degradation and translational repression, however, the measurement of mRNA levels has been shown to be a good representation of miRNA regulation in mammalian cells [20,21]. The miR-205 Tg was also induced with doxycycline for 24 hours, resulting in only 3.5 fold overexpression (Fig. S1c), likely due to the high expression of endogenous miR-205 in keratinocytes [19].

We performed an optimized version of CLEAR-CLIP to enhance quantitativeness and sensitivity (Fig. 1a and see Method). To reduce bias in the ligation steps and enhance ligation efficiency, we used a 5’ linker with a random NNNN and a 3’ linker with a NN at the ligated end of each adaptor and performed the ligation steps in the presence of PEG-8000, respectively. These steps were effective at reducing bias and improving ligation efficiency for small RNA ligation and sequencing [17]. CLEAR-CLIP reads were barcoded using the 5’ NNNN to distinguish unique events, allowing removal of PCR duplicates and in total we sequenced 1,230,019 unique miRNA:RNA chimeras from 18 libraries. Additionally, we used an improved Proteinase K/SDS method to isolate chimeric RNA from the nitrocellulose membrane for improved RNA isolation [22]. To assure robust target detection, we required a mRNA region to be ligated to the same miRNA in at least 2 libraries, which we will hereafter refer to as “high confidence” sites.

**Figure 1.**
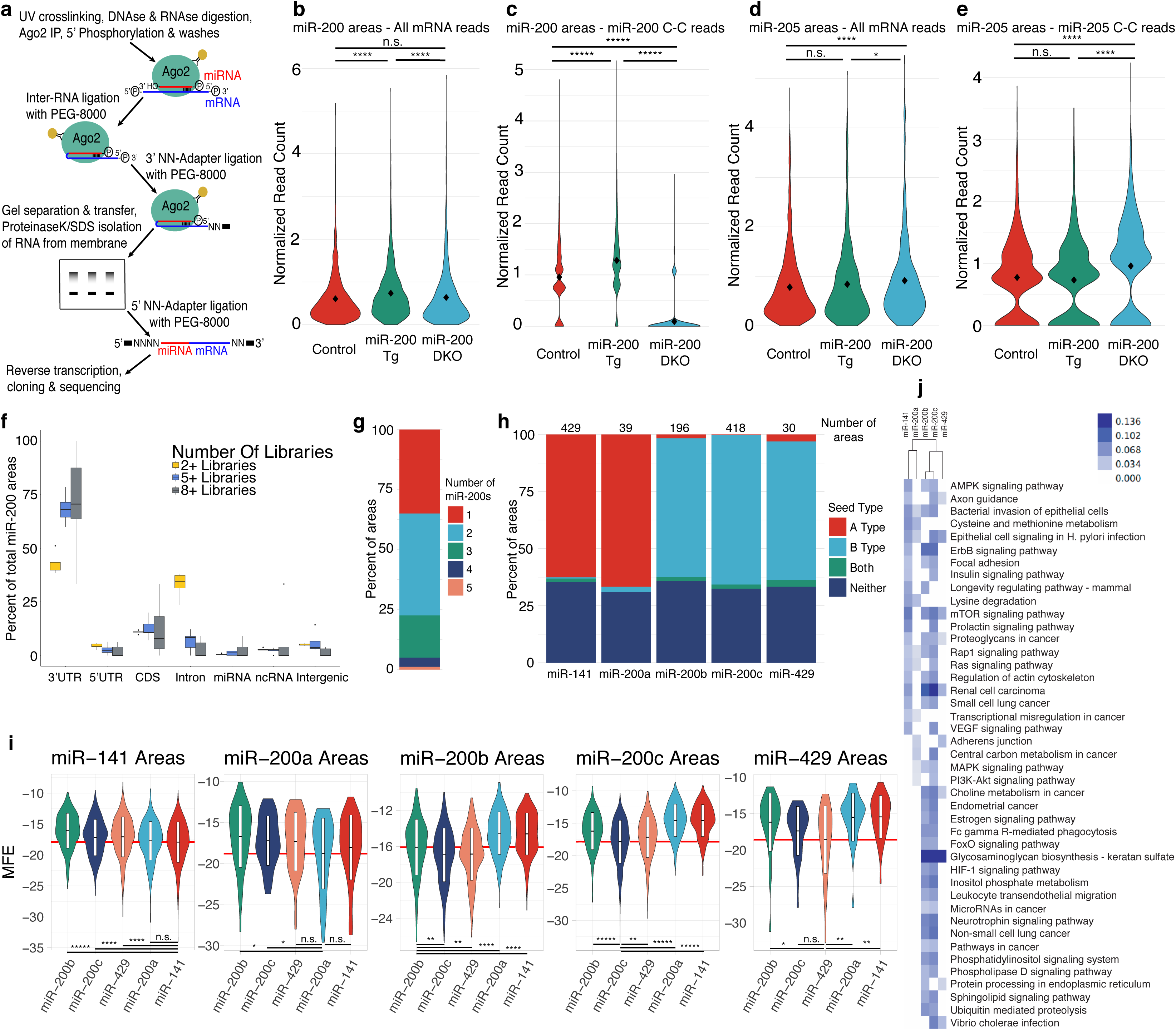
Characterization of miR-200 family binding by CLEAR-CLIP. **a**, A simplified schematic of the CLEAR-CLIP protocol is shown with a focus on changes that we made to the protocol. **b**, The number of mRNA reads in miR-200 high confidence areas is shown as a violin plot for controls, miR-200 Tg and miR-200 DKO samples. **c**, The number of miR-200 specific CLEAR-CLIP reads in miR-200 high confidence areas is shown for controls, miR-200 Tg and miR-200 DKO samples. **d**, The number of mRNA reads in miR-205 high confidence areas is shown for controls, miR-200 Tg and miR-200 DKO samples. **e**, The number of miR-205 specific CLEAR-CLIP reads in miR-205 high confidence areas is shown for controls, miR-200 Tg and miR-200 DKO samples. Statistics a-d: Unpaired two-sided t-test. *, P < 0.05; ****, P < 0.0001; *****, P < 2.2e-16. **f**, Genomic annotations for miR-200 high confidence areas are shown for areas found in 2+, 5+ or 8+ libraries. **g**, The number of miR-200s observed in each high confidence area is shown for all miR-200 high confidence areas. **h**, The percent of areas with a miR-200a type seed, miR-200b type seed, both seeds or neither is displayed for miR-200 areas that had a majority of their reads from one family member. **i**, Binding energy is shown for each miR-200 family member in areas that had the majority of their reads from one family member. The box within each violin plot shows the mean +/− the standard deviation. Unpaired two-sided t-test. *, P < 0.05; **, P < 0.01; ****, P < 0.0001; *****, P < 2.2e-16. **j**, The combined scores from pathway enrichment for miR-200 family members were subjected to hierarchical clustering and displayed as a heat map.

For the miR-200 family, we allowed chimeric reads with any miR-200 family member found in 2 or more libraries (both control and inducible samples) to define a high confidence site, which resulted in 2,369 miR-200 sites in 3’UTRs, corresponding to 1,486 unique genes. We first compared the sensitivity of detecting miRNA targeted sites between mRNA only reads that are typically obtained in HITS-CLIP and miRNA:mRNA chimeric reads that are obtained by CLEAR-CLIP. When we examined the relative read density of these high confidence sites using mRNA only fragments, we observed little to no difference between control, miR-200 Tg and DKO samples (Fig. 1b). In contrast, when we used miR-200:mRNA chimeric reads to specifically examine miR-200-mediated targeting events, we observed an increase in miR-200 Tg samples and an almost total loss in the DKO samples as expected (Fig. 1c). As a control, we also calculated read density within the high confidence sites of miR-205, a highly expressed but unrelated miRNA, in control, miR-200 Tg and DKO samples. We did not observe any change in mRNA coverage for these miR-205 sites between control and miR-200 Tg and a slight increase in the miR-200 DKO when using mRNA only reads (Fig. 1d). When miR-205:mRNA chimeric reads were examined, there was again no change between control and miR-200 Tg but a larger increase in the miR-200 DKO sample, reflecting a higher sensitivity and quantitative performance of CLEAR-CLIP (Fig. 1e). We also compared our CLEAR-CLIP data to HITS-CLIP previously published by our lab [23] and found that for both miR-200s and miR-205, CLEAR-CLIP identified more miRNA targets that resulted in better gene repression (Figs. S1d-e). Together, these data indicate the high sensitivity and specificity of CLEAR-CLIP for identifying miRNA-interacting sites in comparison to HITS-CLIP. Furthermore, these results suggest that in the absence of miR-200s there is an increase in targeting by other miRNAs.

We next examined what regions of the genome the mRNA portion of miR-200 high confidence sites annotated to. When requiring areas to be found in 2 or more libraries, ∼45% of sites annotated to 3’UTRs, 35% to introns and small percentages to 5’UTRs, CDS, miRNAs, non-coding RNAs (ncRNAs) and intergenic regions (Fig. 1f). However, ∼70% of sites annotated to 3’UTRs and ∼10% annotated to introns when requiring sites to be found in 5+ or 8+ libraries (Fig. 1f). These results indicate that even though chimeras to intronic regions are seen with some frequency, reproducible sites are usually found in 3’UTRs. Additionally, to determine whether miR-200 sites from different regions of the genome were functional, we selected genes that had only one miR-200 high confidence site and analyzed their expression when miR-200s were induced. We found 3’UTR sites to be highly effective, while CDS sites had a slight but statistically significant effect and other regions including 5’UTR and intron were not effective for gene repression (Fig. S1f). We subsequently focused our studies on 3’UTR sites.

Because miR-200 miRNAs from the two sub-families differ by one nucleotide at the fourth position in their seed sequences, we inspected how many individual miR-200 family members recognize the same site. The majority of miR-200 sites were targeted by one (35% of sites) or two (42.5% of sites) family members, but some sites were targeted by 3 (17.5%), 4 (3.8%) or all 5 (1.2%) family members (Fig. 1g). The majority of the sites (80.7%) had reads from the same seed family, while the rest (19.3%) had reads from both miR-200 seed sub-families (Fig. S1g). We next analyzed how often each miR-200 member bound its cognate seed versus the opposite seed. Overall, ∼60% of the sites for all miR-200 family members contained the cognate seed and ∼30% did not contain either seed (Fig. 1h). A small percentage of sites contained either both seeds or the opposite seed, indicating that miR-200 family members are much more likely to bind a site with the cognate seed even though their seed sequences only differ by one nucleotide. Furthermore, to examine the specificity of each family member, RNAhybrid [24] was used to calculate the predicted binding energy for all five family members against sites that were found to be dominated by an individual family member. Most sites that had a majority of reads from one family member correspondingly had the lowest binding energy for that member, and seed families also had lower binding energy within their family (Fig. 1i). Interestingly, miR-200c and miR-429 had lower average binding energies to miR-200b sites than miR-200b itself, likely due to the lower binding energy of miR-200c and miR-429 to their perfect reverse complementary target than miR-200b (−43.3 kcal/mol and −40.3 kcal/mol, respectively versus −38.5 kcal/mol for miR-200b).

To probe the genes regulated by this five-member family, we performed pathway analysis on their high confidence targets. Using high confidence sites with a seed (7mer or better) in 3’UTRs we compiled a gene list for each miR-200 family member. Gene Ontology (GO) was then performed on these gene lists examining for enrichment of KEGG pathways using Enrichr [25]. Focal adhesion, Ras signaling and PI3K-Akt signaling pathways were found to be regulated by more than one family member using the hierarchical clustering of GO terms (Fig. 1j and Fig. S1h). Interestingly, many GO categories were targeted by both seed families, supporting a coordinated targeting mechanism by the miR-200 family [18].

### CLEAR-CLIP identifies functional miRNA targeting sites

miRNA levels often dramatically change during homeostasis such as cell lineage specification, during stress responses such as wound healing as well as under pathological conditions such as tumorigenesis [4,26]. Furthermore, many studies have relied on overexpression to examine miRNA functions. However, it is unknown whether elevated miRNA expression preferentially represses existing targets or inhibits new targets [9]. To address this issue, we examined whether overexpression of the miR-200b/a/429 cluster above physiological levels results in off-target effects, causing miRNAs to target genes not seen at physiological miRNA levels. We quantified the normalized read numbers in miR-200 high confidence sites between control and induced samples. We noticed there was a shift towards higher relative reads numbers in induced samples, as expected, and we also found a number of sites that were found only in the control or only in the induced (Fig. 2a). We also examined repression of these targets upon induction of the miR-200b cluster. Genes that contain the miR-200 sites in both control and induced samples (1,091 genes) were better repressed by miR-200 induction than genes containing sites seen only in control (98 genes) or induced (252 genes) samples (Fig. 2b). The stronger repression of shared sites was consistent with the observation that these sites had more CLEAR-CLIP reads when more miR-200 were expressed in induced samples (Fig. 2c). These data indicate that elevated expression of miRNAs (∼15-fold) primarily results in heavier targeting of canonical sites.

**Figure 2.**
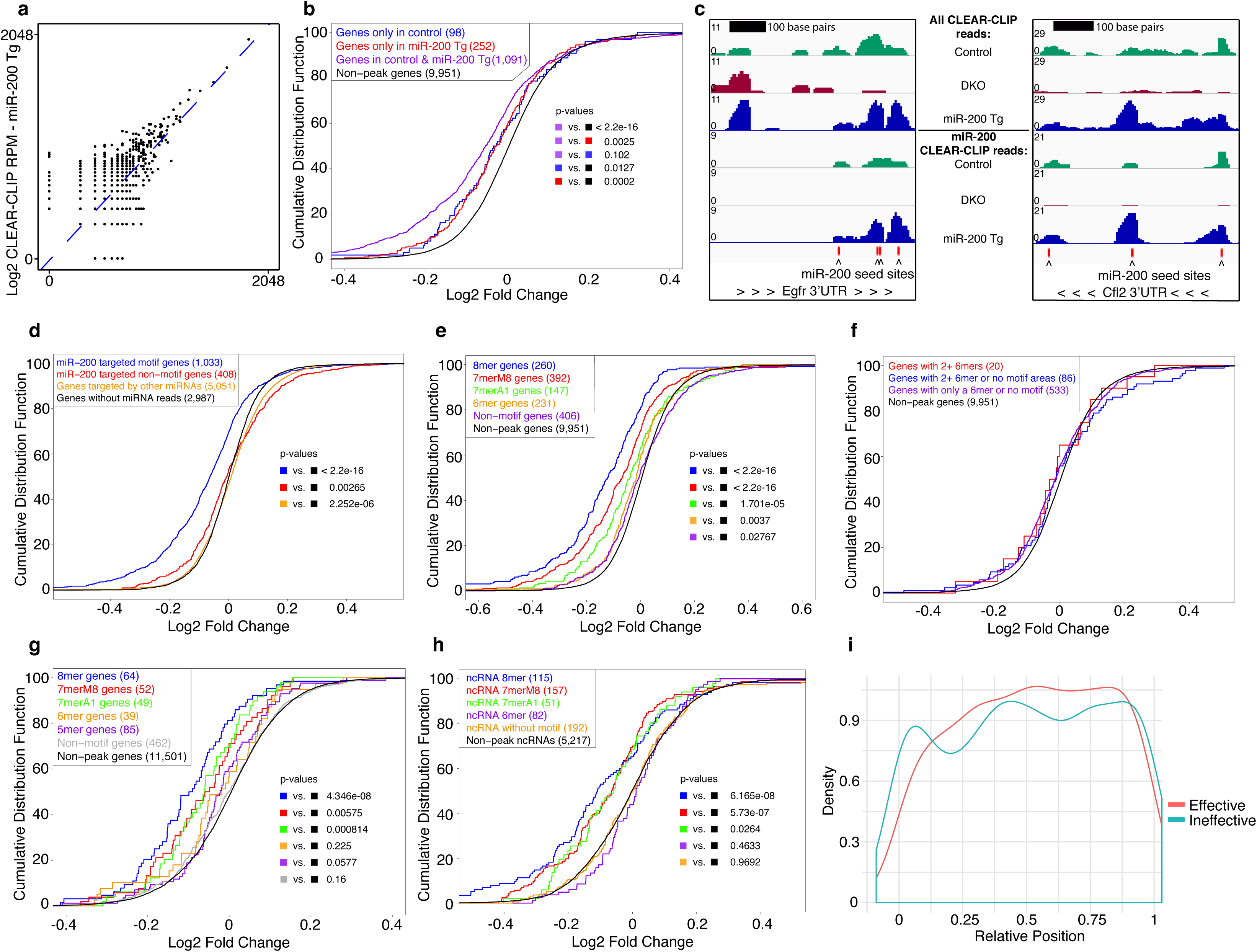
Characterization of miR-200 and miR-205 regulation of target genes. **a**, Correlation of CLEAR-CLIP reads per miR-200 high confidence site between controls and miR-200 Tg. Plotted as Log2 Reads Per Million (RPM) mapped CLEAR-CLIP reads. **b**, Log2 fold change in gene expression upon induction of the miR-200b cluster on genes found by CLEAR-CLIP in controls, miR-200 Tg or both. **c**, Example tracks from Egfr and Cfl2 are shown for CLEAR-CLIP reads from all miRNAs (top 3 tracks) and miR-200 CLEAR-CLIP reads (bottom 3 tracks). Controls, miR-200 DKO and miR-200 Tg are shown separately. Scale is denoted on the left for each track. miR-200 seed sites are denoted on the bottom with red boxes. **d**, Log2 fold change in gene expression upon induction of the miR-200b cluster is shown for miR-200 motif genes versus genes without a miR-200 motif, genes targeted by miRNAs other than miR-200 and genes without any CLEAR-CLIP reads. **e**, Log2 fold change in gene expression upon induction of the miR-200b cluster is shown for genes with canonical motifs, versus without a motif and genes without a high confidence miR-200 peak. **f**, Log2 fold change in gene expression upon induction of the miR-200b cluster is shown for genes with multiple 6mer motif or non-motif areas as compared to non-peak genes. **g**, Log2 fold change in gene expression upon induction of miR-205 is shown for canonical motifs and genes with a 5mer motif (nucleotides 3-7) as compared to non-motif and non-miR-205-peak genes. **h**, Log2 fold change in gene expression upon induction of the miR-200b cluster is shown for poly(A) selected ncRNAs with canonical miR-200 motifs as compared to poly(A) selected ncRNAs without a high confidence miR-200 peak. **i**, The center of miR-200 CLEAR-CLIP peaks is plotted along the relative length of ncRNAs that were observed to be repressed by miR-200 induction (Effective) or not (Ineffective). For all CDF plots the number of genes is shown in parenthesis and p-values were calculated using the Kolmogorov–Smirnov test.

Because ∼30% of miR-200 CLEAR-CLIP areas did not contain a seed match (Fig. 1h), we next examined the effectiveness of CLEAR-CLIP identified target sites in mediating gene repression. Genes with a miR-200 high confidence site with or without a miR-200 seed (6mer or better) were both significantly repressed by induction of miR-200s. However, genes with the seed motif were repressed significantly better than genes without a motif (Fig. 2d). Interestingly, genes containing CLEAR-CLIP identified target sites of other miRNAs were slightly derepressed upon induction of miR-200s, indicating competition for the availability of the RISC by induced miR-200s[27]. Additionally, genes that were derepressed (> 0.05 log2 fold change) tended to have lower expression level (Fig. S2a) and more total CLEAR-CLIP reads (Fig. S2b) as compared to genes that were not derepressed.

To further analyze the impact of different types of seed matches, we classified miR-200 targets into categories by the best miRNA motif they contained and analyzed their effectiveness in mediating gene repression (Fig. 2e). Targets containing 8mer, 7merM8 and 7merA1 motifs were well repressed with the stronger match conferring stronger inhibition, consistent with previous reports [2]. Statistically significant repression was also seen for genes with a 6mer match or without any seed match, but the repression for both groups of genes was much weaker (Fig. 2e). Furthermore, genes with multiple miR-200 CLEAR-CLIP sites harboring a 6mer or without any seed match were not repressed more than genes with a single 6mer or no motif, reflecting general ineffectiveness of these sites (Fig. 2f). These data suggest that, at least for miR-200s, a 7mer or 8mer match in a CLEAR-CLIP identified site is critical for effective repression.

Similarly, the effectiveness of motifs was examined in miR-205 high confidence sites. Compared to miR-200s, miR-205 sites showed fewer canonical 7mer or 8mer motifs and more sites that did not have a seed match. Therefore, we performed *de novo* motif analysis using HOMER [28] on miR-205 sites that did not have a canonical seed match. This analysis found a 5mer match (nucleotides 3 to 7 of the miRNA) that was prevalent in 22% of the non-seed matching sites (Fig. S2c). When we combined miR-205 CLEAR-CLIP identified sites with miR-205 Tg RNA-Seq, we found the dependence on miR-205 seed matches less apparent than for miR-200s and only observed significant repression of genes that contained an 8mer or 7mer match (Fig. 2g). These data suggest that the effectiveness of a seed match may vary among different miRNAs.

We next examined whether miR-200s could be targeting non-coding RNAs (ncRNAs) that are annotated in a database of murine ncRNAs[29]. We found 1,292 miR-200 high confidence sites in 806 unique ncRNA genes. Among these, 597 ncRNA genes were detected in our poly(A) selected RNA-Seq data. Notably, ncRNAs with miR-200 high confidence sites containing an 8mer or 7mer were significantly repressed (Fig. 2h), similar to coding genes. This suggests that miR-200s may also play a role in repressing ncRNAs. Because of the 3’UTR bias of effective miRNA targeting for coding genes, we examined whether miR-200s were targeting a particular region on ncRNAs. The center of miR-200 high confidence sites was plotted along the length of the ncRNA (normalized to 1) for ncRNAs that were repressed by more than 0.1 log2 fold change (effective targets) in our miR-200 Tg RNA-Seq and compared to ncRNAs that were not repressed (ineffective targets). No positional bias of miR-200 target sites along the length of ncRNAs was observed for either effective or ineffective sites (Fig. 2i). Because this result could be confounded by the length of the ncRNA, we further examined the distance of miR-200 sites to the 3’ end of ncRNAs and did not observe a bias of effective sites towards the 3’ end (Fig. S2e).

### Different miRNAs have a different degree of reliance on the seed match

Because the miR-200 family harbors two distinct seed sequences and these five miRNAs are co-expressed in epithelial cells, we assessed whether repression of miR-200 high confidence targets was affected by containing one or both seed types. Genes that contained both a miR-200a and miR-200b type seed were repressed better upon miR-200 induction than genes that contained either one miR-200a type or miR-200b type seed (Fig. 3a). Genes with both seed types were repressed similarly to genes with two miR-200b type seeds, but significantly better than genes with two miR-200a type seeds. This may be due to the fact that we induced expression of the miR-200b cluster that expresses two miR-200b type miRNAs and only one miR-200a type miRNA. We next examined whether genes that are targeted by more members of the miR-200 family are better repressed by induction of miR-200s. Indeed, the more miR-200 family members that were found to be associated with a gene by CLEAR-CLIP, the better repression upon miR-200 induction (Fig. 3b).

**Figure 3.**
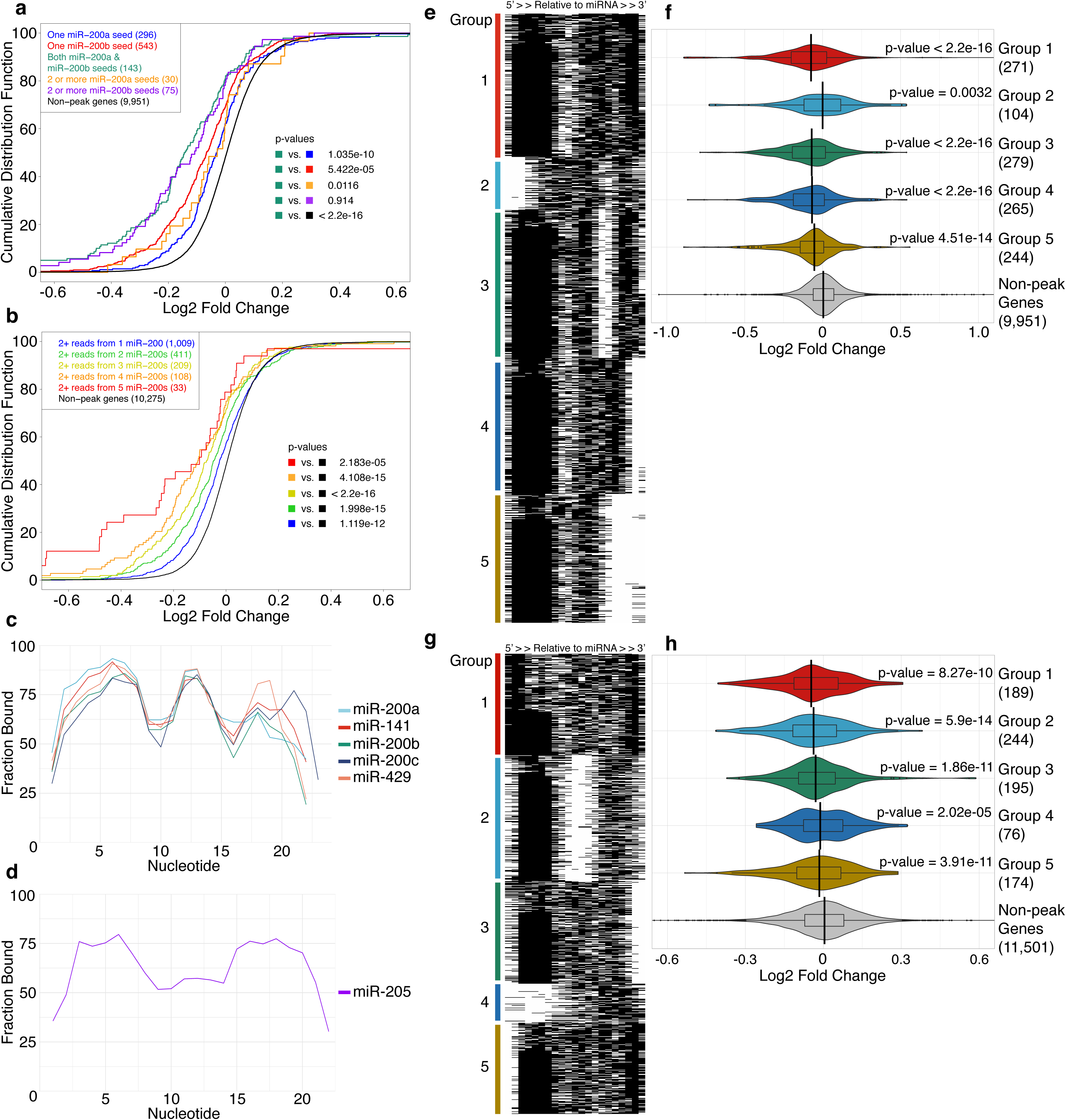
miR-200s are highly dependent on seed matches for target repression. **a**, Log2 fold change in gene expression upon induction of the miR-200b cluster is shown for genes that contain one miR-200a type seed, one miR-200b type seed, both seed types or 2 or more of each seed type. **b**, Log2 fold change in gene expression upon induction of the miR-200b cluster is shown for genes that contain 2 or more reads from 1, 2, 3, 4 or 5 miR-200s as compared to non-miR-200 peak genes. **c**, The predicted fraction bound is displayed for each nucleotide along the length of each miR-200 for all of that member’s high confidence areas. **d**, The predicted fraction bound is displayed for each nucleotide along the length of miR-205. **e**, High confidence miR-200 areas with the majority of reads for that area from a single family member were hybridized using RNAhybrid and the individual nucleotides were predicted to be bound or not bound. All miR-200 family members were then pooled and the predicted nucleotide hybridization was clustered in 5 groups using k-means clustering. This hybridization was then graphed as a heat map with black denoting the nucleotide is bound and white meaning not bound. **f**, Log2 fold change in gene expression upon induction of the miR-200b cluster is shown for the 5 k-means clusters from e, as compared to non-miR-200 peak genes. The indicated p-value for each is compared to non-miR-200 peak genes. **g**, High confidence miR-205 areas were hybridized against miR-205 using RNAhybrid, with the seed being enforced if one was present within the area. Predicted binding was then clustered into 5 k-means clusters and graphed as a heat map with black denoting the nucleotide is bound and white meaning not bound. The indicated p-value for each is compared to non-miR-205 peak genes. **h**, Log2 fold change in gene expression upon induction of miR-205 is shown for the 5 k-means clusters from g, as compared to non-miR-205 peak genes. For all CDF and violin plots the number of genes is shown in parenthesis and p-values were calculated using the Kolmogorov–Smirnov test.

To further characterize binding by miR-200s, high confidence sites for each miR-200 member were generated individually and RNAhybrid was used to calculate the best binding site within each area. This information was then used to calculate how often each nucleotide was paired to its mRNA target. As shown in figure 3c, miR-200s have a strong preference for a seed match. Interestingly, a large percentage of miR-200 areas also used nucleotides 12-14 of the miRNA for binding their targets. The binding fraction by nucleotide was also calculated for miR-205 (Fig. 3d). By comparison, miR-205 depends more heavily on nucleotides 3-7 of its seed, and utilizes more 3’ end binding than nucleotides 12-14. These data suggest that 3’ end binding is also variable among different miRNAs. Additionally, the predicted binding from RNAhybrid was used to classify different binding subcategories by k-means clustering all binding subsets of the miR-200 family members and then examining their ability to repress gene expression. As shown in figure 3e, miR-200s clustered into four mostly seed containing groups with different modes of 3’ end binding (groups 1,3,4,5) and one group that lacked seed matches (group 2). Examining gene expression upon the induction of miR-200s, it was evident that the four groups with a seed match resulted in similar repression, whereas group 2, which lacked a seed match, resulted in less repression (Fig. 3f). We also performed the same analysis for miR-205 high confidence targets (Fig. 3g) and again observed more variability in miR-205 binding using its seed region. Assaying the functionality of these groups using RNA-seq revealed groups 1, 2, 3 and 5, which contain both seed and 3’ end matches, were significantly repressed. Group 4 genes, which did not contain seed matches, were also significantly repressed, but less so (Fig. 3h). These data show that miR-200s are highly dependent on a seed match, whereas miR-205 is more dependent on both seed and 3’ end binding.

### Quantitative analysis of CLEAR-CLIP identified miRNA targets

We next tested whether our optimized CLEAR-CLIP can predict the strength of miR-200-dependent regulation based on the number of captured miRNA:RNA chimeric reads. We first examined the correlation between the number of miR-200 CLEAR-CLIP reads per gene and their repression upon miR-200 induction. Genes with increasing numbers of CLEAR-CLIP reads were more repressed upon induction with miR-200s (Fig. 4a). Additionally, when miR-200 chimeric reads count for 20% or more of total CLEAR-CLIP reads, they also confer stronger repression by miR-200 induction (Fig. 4b). We further binned miR-200 targets into three groups based on reads per gene (1-6, 7-30 or 31+ reads) and further divided each group into two categories: low percent miR-200 targeting (< 20%) or high percent miR-200 targeting (> 20%). Interestingly, high percentage miR-200 targeting was correlated with better gene repression for each group (Fig. 4c). Similarly, for miR-205, more miR-205 CLEAR-CLIP reads per gene was also correlated with stronger repression. Genes with 3-9 or 10+ reads were significantly repressed by miR-205 induction, in contrast to genes with only 1-2 reads (Fig. 4d). Next, we also examined miR-205 repression by the percent of miR-205 reads out of total CLEAR-CLIP reads per gene. We found that genes where miR-205 constituted >10% of the total CLEAR-CLIP reads were repressed whereas genes where miR-205 constituted 0-10% of the total reads were not significantly repressed (Fig. 4e). Again, when miR-205 targets were binned into three groups based on reads per gene (1-2 reads, 3-9 reads and 10+ reads) and then further divided into high and low percent miR-205 targeting (more or less than 10%), we found for each group the set of genes with a high percent of miR-205 reads was repressed better than the low percentage set (Fig. 4f). Finally, we also observed a strong correlation between the number of discrete CLEAR-CLIP sites per gene, especially for genes with > 3 sites and the number of sites with a seed match per gene on repression upon the induction of miR-200 (Fig. S3a-b) or miR-205 (Fig. S3c). Together, these analyses show that optimized CLEAR-CLIP quantitatively reflects the strength of miRNA-mediated regulation.

**Figure 4.**
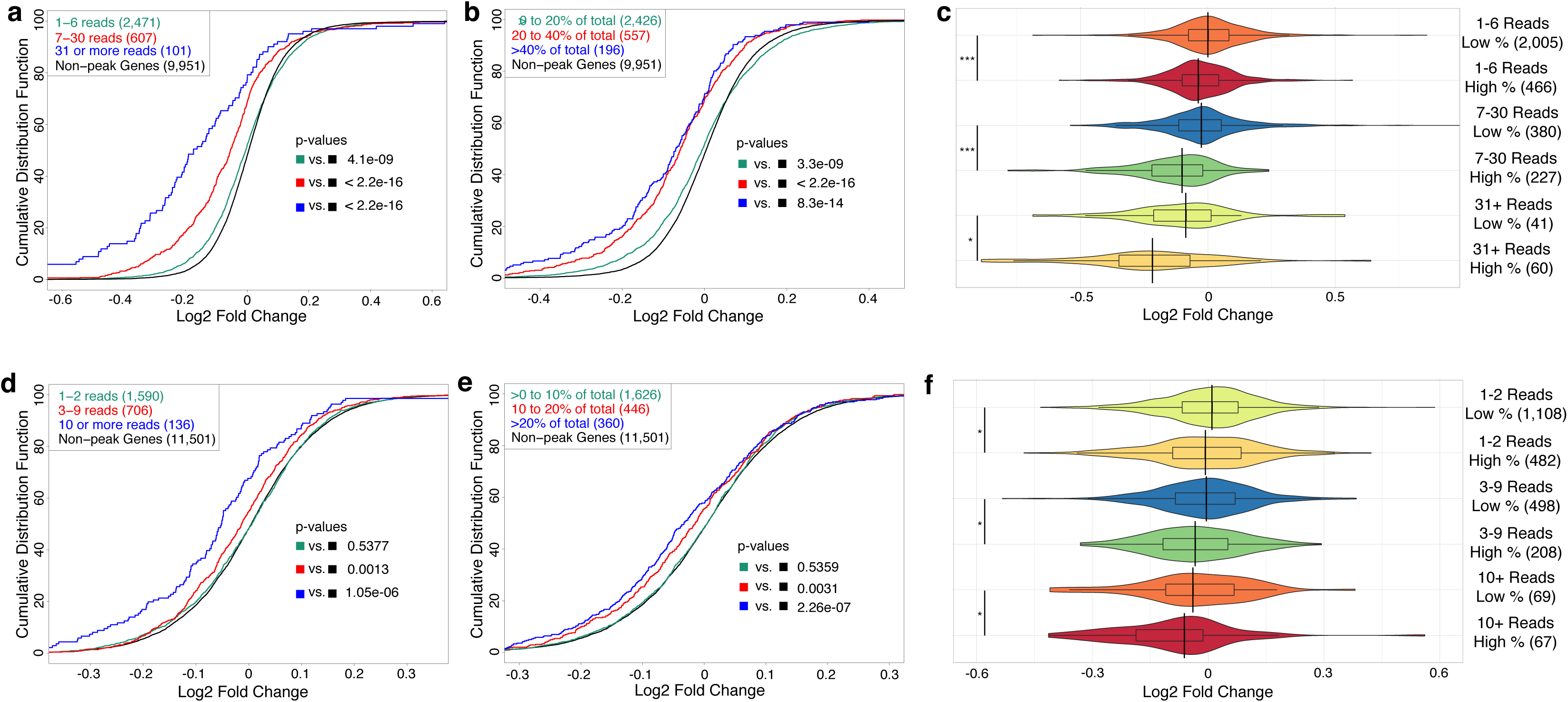
Quantitative capture of miRNA:RNA interactions predicts the strength of miRNA-mediated repression. **a**, Log2 fold change in gene expression upon induction of the miR-200b cluster given the number of miR-200 family reads per gene as compared to non-miR-200 high confidence genes. **b**, Log2 fold change in gene expression upon induction of the miR-200b cluster given the percentage of miR-200 reads out of total CLEAR-CLIP reads per gene. **c**, Log2 fold change in gene expression upon induction of the miR-200b cluster given a low or high percentage of miR-200 reads out of total CLEAR-CLIP reads, binned into groups of 1-6 reads, 7-30 reads and 31+ reads. **d**, Log2 fold change in gene expression upon induction of miR-205 given the number of miR-205 reads per gene as compared to genes that did not have a miR-205 high confidence area. **e**, Log2 fold change in gene expression upon induction of miR-205 given the percentage of miR-205 reads out of total CLEAR-CLIP reads per gene. **f**, Log2 fold change in gene expression upon induction of miR-205 given a low or high percentage of miR-205 reads out of total CLEAR-CLIP reads, binned into groups of 1-2 reads, 3-9 reads and 10+ reads. For all panels the number of genes in each category is shown in parenthesis. *, P < 0.05; *** P < 0.0001. All statistics for figure calculated using the Kolmogorov–Smirnov test.

### Comparison between CLEAR-CLIP identified targets and TargetScan predicted targets

CLEAR-CLIP and computational algorithms such as TargetScan are two different approaches that can provide miRNA- and site-specific information for miRNA targeting. We therefore compared the performance of our CLEAR-CLIP method and the latest TargetScan predictions for mouse (TargetScan 7.1 evolutionarily conserved miRNA sites). Because CLEAR-CLIP identifies targets which are expressed in a cellular context and are bound by miRNAs regardless whether they harbor seed matches, and TargetScan predicts all targets irrespective of gene expression, we required genes of interest to be expressed in our system (basemean > 10 in our RNA-Seq) and required our CLEAR-CLIP targets to have a 7mer or 8mer seed match, identical to target lists we used from TargetScan. For both miR-200 seed types, we identified 436 expressed genes that were shared between our CLEAR-CLIP data and TargetScan predictions (Fig. 5a). We also found 366 genes that were only identified in our CLEAR-CLIP and 854 genes that were only predicated by TargetScan. To determine the effectiveness of these targets regulated by miR-200s, we determined the repression of CLEAR-CLIP targets versus TargetScan predictions upon miR-200 induction. Overall, CLEAR-CLIP targets were better repressed upon miR-200 induction than TargetScan predictions (Fig. 5b). Furthermore, genes found both by CLEAR-CLIP and predicted by TargetScan were best repressed by miR-200 induction, followed by genes only identified by CLEAR-CLIP and then genes only predicted by TargetScan (Fig. 5c).

**Figure 5.**
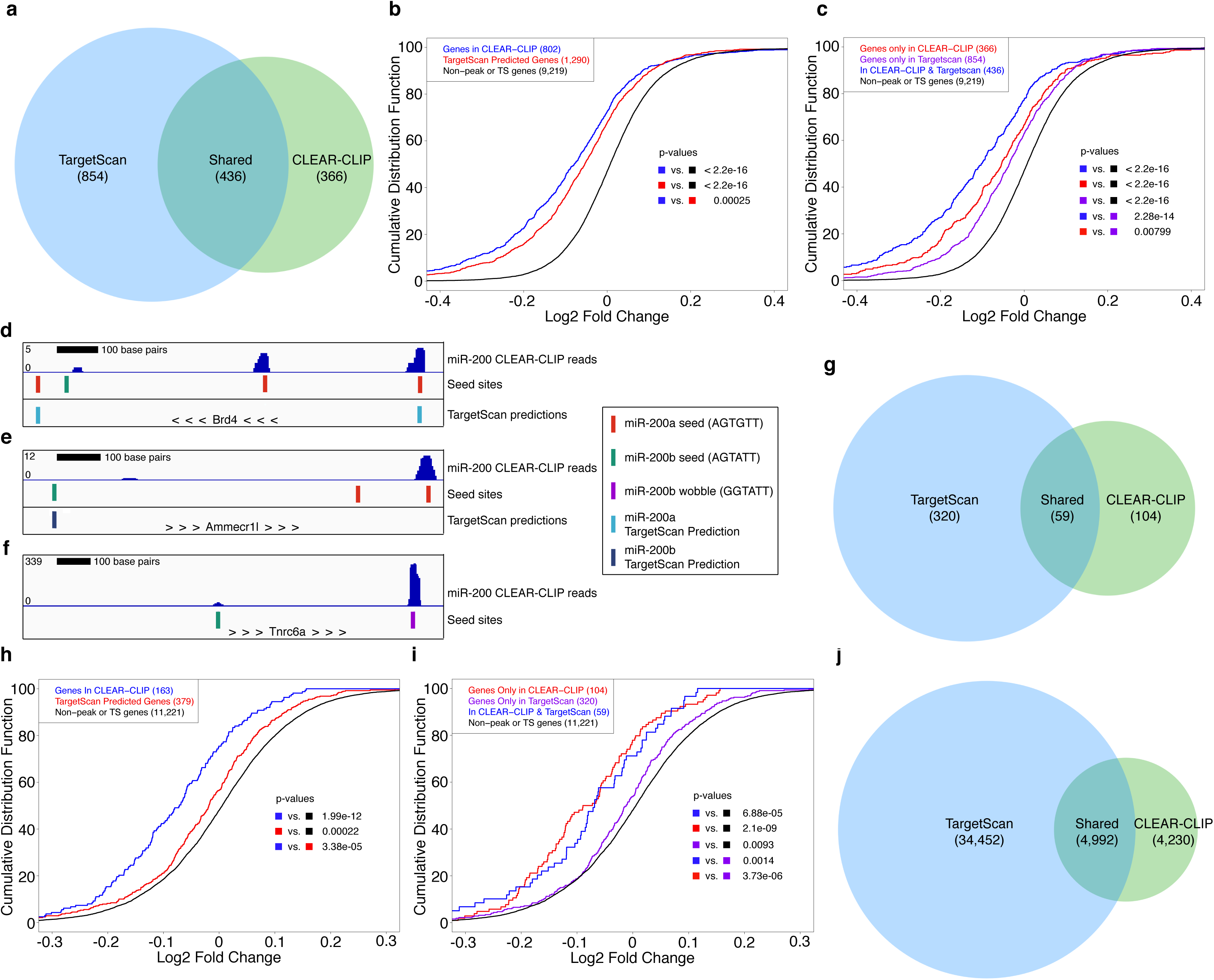
Comparison of performance between CLEAR-CLIP captured and TargetScan predicted targets. **a**, Overlap between miR-200 CLEAR-CLIP genes with a 7mer or 8mer and TargetScan predictions for miR-200s. **b**, Log2 fold change in gene expression upon induction of the miR-200b cluster is shown for CLEAR-CLIP genes with a 7mer or 8mer and TargetScan predicted conserved sites as compared to genes without a miR-200 high confidence site and not predicted as conserved by TargetScan. **c**, Log2 fold change in gene expression upon induction of the miR-200b cluster is shown for genes only in CLEAR-CLIP, only predicted by TargetScan or in CLEAR-CLIP and TargetScan as compared to genes without a miR-200 high confidence site and not predicted by TargetScan. **d**, A portion of the Brd4 3’UTR is shown with miR-200 CLEAR-CLIP reads (top track), miR-200 seed sites (middle track) and TargetScan sites (bottom track) indicated. **e**, A portion of the Ammecr1l 3’UTR is shown with miR-200 CLEAR-CLIP reads (top track), miR-200 seed sites (middle track) and TargetScan sites (bottom track) indicated. **f**, A portion of the Tnrc6a 3’UTR is shown with miR-200 CLEAR-CLIP reads (top track) and miR-200 seed sites (bottom track) indicated. **g**, Overlap between miR-205 CLEAR-CLIP genes with a 7mer or 8mer and TargetScan predicted conserved sites for miR-205. **h**, Log2 fold change in gene expression upon induction of miR-205 is shown for CLEAR-CLIP genes with a 7mer or 8mer and TargetScan predicted sites as compared to genes without a miR-205 high confidence site and not predicted as conserved by TargetScan. **i**, Log2 fold change in gene expression upon induction of miR-205 is shown for genes only in CLEAR-CLIP, only predicted by TargetScan or in CLEAR-CLIP and TargetScan as compared to genes without a miR-205 high confidence site and not predicted by TargetScan. **j**, Overlap between all miRNA:mRNA CLEAR-CLIP interactions with a 7mer or 8mer and all conserved TargetScan predictions. For all CDF plots the number of genes is shown in parenthesis and p-values were calculated using the Kolmogorov–Smirnov test.

To determine whether gene expression contributes to the lack of detection by CLEAR-CLIP, we examined the expression level in our RNA-Seq data of genes found in the different overlaps. Genes predicted only by TargetScan and not found in our CLEAR-CLIP did have slightly lower average expression level, but many of the genes only predicted by TargetScan were expressed at a similar level to genes detected by CLEAR-CLIP (Fig. S3d). These data suggest that the lack of detection was not simply due to the low expression. Furthermore, although many common targets between CLEAR-CLIP and TargetScan prediction were indeed based on the same sites, sometimes a gene was found by CLEAR-CLIP and TargetScan, but targeted at different sites along the 3’UTR. For example, Brd4 contained one site that was captured by CLEAR-CLIP and also predicted by TargetScan, but it also had one site with a seed only captured by CLEAR-CLIP, and a TargetScan predicted site that did not have any CLEAR-CLIP reads (Fig. 5d). Further, Ammecr1l contained one robust CLEAR-CLIP site with a seed that was not predicted by TargetScan and a predicted site that did not have any miR-200 CLEAR-CLIP reads (Fig. 5e). Additionally, TargetScan misses any sites that lack a canonical seed. For example, we found a heavily targeted site in the 3’UTR of Tnrc6a that lacked a canonical seed and instead had a match with a G:U wobble (GGTATT instead of AGTATT) (Fig. 5f). Tnrc6a was also down 20% upon induction of miR-200.

To extend the study beyond miR-200s, we performed a similar comparison for targets of miR-205 captured by CLEAR-CLIP and predicted by TargetScan. Possibly due to a lower percentage of miR-205 sites containing a 7mer or 8mer seed, there was even less overlap between CLEAR-CLIP and TargetScan predications. We found only 59 genes identified in both. TargetScan predicted 320 genes that were not identified by CLEAR-CLIP and our CLEAR-CLIP identified 104 genes that were not predicted by TargetScan (Fig. 5g). Despite identifying fewer targets, we still found that CLEAR-CLIP targets were repressed more effectively than TargetScan predications upon induction of miR-205 (Fig. 5h). Furthermore, the 104 genes only detected by CLEAR-CLIP were repressed similarly to the 59 common targets. The 320 genes only predicted by TargetScan were minimally repressed (Fig. 5i).

Next, we performed a global comparison of CLEAR-CLIP and TargetScan predictions. We used all miRNAs with at least 50 high confidence target sites and analyzed their predicted versus captured sites. High confidence miRNA:RNA interactions from our data were defined as those that are found in 2+ libraries, in 3’UTRs, with an mRNA basemean 10+ in our RNA-Seq and with a 7mer or better seed. We found 4,992 miRNA:RNA interactions that were shared between our CLEAR-CLIP and TargetScan predictions. However, 34,452 interactions were predicted by TargetScan but not captured by CLEAR-CLIP and 4,230 interactions were captured by CLEAR-CLIP but not predicted by TargetScan (Fig. 5j). These results indicate the differences in the methods and highlight the value of direct identification of targets for narrowing down miRNA recognized genes over prediction algorithms.

### CLEAR-CLIP allows genome-wide discovery of miRNA-regulated gene networks

We next examined global features of miRNA-mediated target recognition by examining well expressed miRNAs. We studied 88 miRNAs that had 50 or more high confidence target sites. First, we calculated high confidence sites for each miRNA that were found in 2+, 5+ or 8+ libraries and then annotated these areas to the genome. Similar to the pattern of miR-200s (Fig. 1f), miRNA target sites are highly enriched in 3’UTRs (Fig. 6a). To map the global binding preference along the length of these 88 miRNAs, we calculated the fractional binding for each nucleotide using RNAhybrid across all high confidence sites. The majority of miRNAs appear to rely heavily on seed binding, but the dependence on seed versus 3’ certainly varies by miRNA (Fig. 6b). Across these miRNAs, the seed nucleotides (2-8) are bound in more than 75% of occurrences. The fraction bound decreases at nucleotides 9 and 10, then increases for nucleotides 11 through 15 and trails off towards the 3’ end of the miRNA (Fig. 6c).

**Figure 6.**
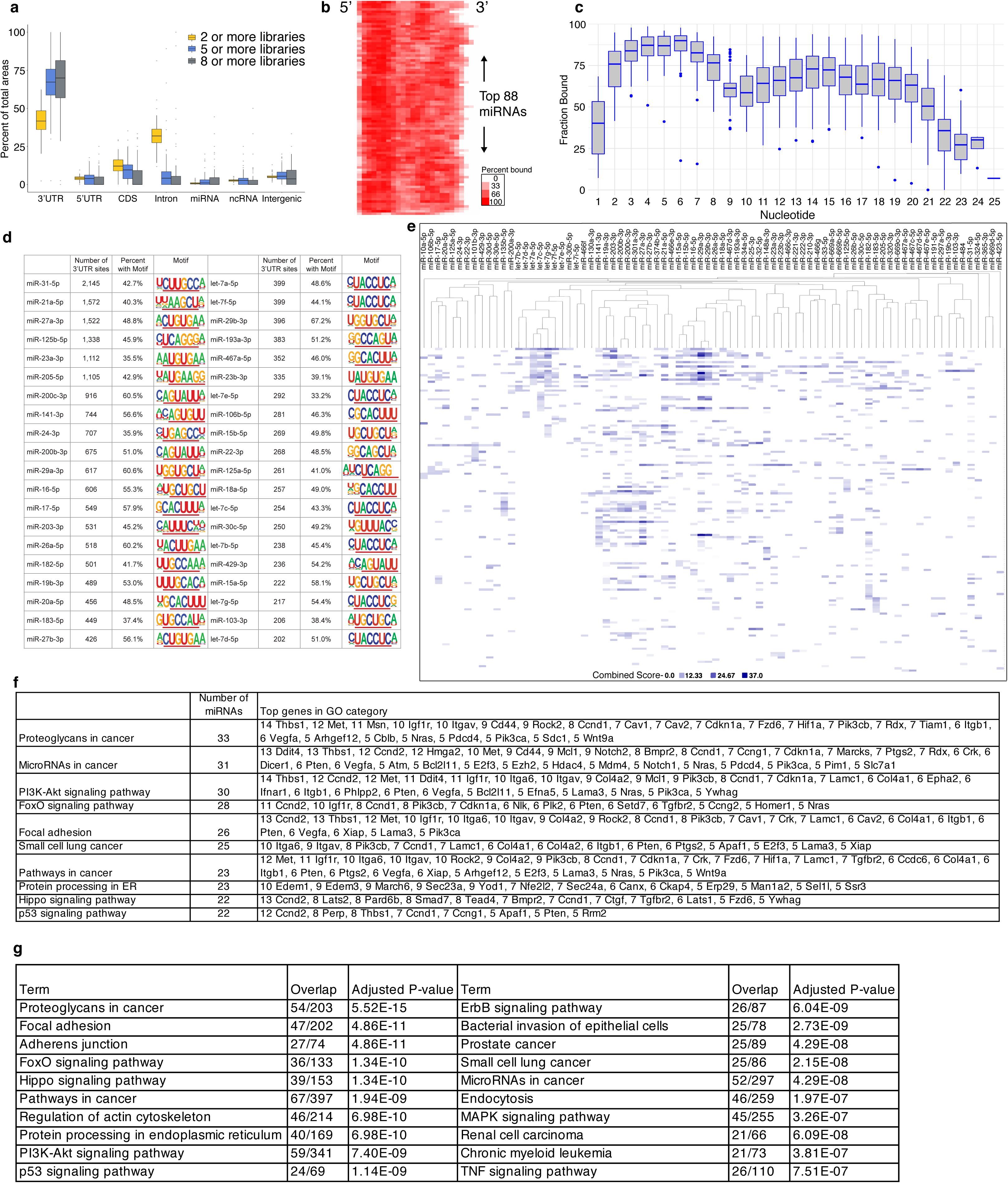
Global analysis of miRNA targeting. **a**, Areas from all miRNAs with 50 or more high confidence sites were annotated to the genome and compared to areas found in 5 or more libraries and 8 or more libraries. **b**, Predicted binding by RNAhybrid is shown for all high confidence areas for all well-expressed miRNAs as a heat map, with darker red indicating that nucleotide of the miRNA is more likely to be bound. **c**, Same data as for (b), except displayed as a box plot of fractional binding along the length of all well-expressed miRNAs. **d**, 3’UTR high confidence areas for the top 40 miRNAs by number of sites were processed by HOMER to detect enriched 8mer motifs. The number of 3’UTR areas observed, the percent with the motif and the motif is shown for each miRNA. **e**, Pathway enrichment was performed for well-expressed miRNAs and the combined score for each enriched pathway was used as a readout to hierarchical cluster miRNAs by what pathways they target. **f**, Top enriched categories from d are shown with the number of miRNAs that were found to have that category enriched and the top genes in that category and how many miRNAs were observed to target them. **g**, Genes that were heavily targeted in our CLEAR-CLIP analysis were used to look for enriched pathways. The top categories are shown here with the number of overlapping genes found and the adjusted p-value.

We next examined the presence of seed matches within captured mRNA target sites of the top 40 miRNAs. For these miRNAs, we searched for enriched 8 nucleotide sequences in high confidence sites in 3’UTRs using HOMER [28]. For most miRNAs, the perfect matches to 6mer seed sequences are the most enriched motif (Fig. 6d). However, a few miRNAs also have some slight variation of the 6mer sequences mostly corresponding to nucleotides 2 or 7, particularly miR-21-5p, miR-203-3p and miR-125a-5p. In addition, the most prolific miRNA, miR-31-5p, binds to 2,145 sites whereas the least prolific miRNA among the top 40 miRNAs, let-7d-5p, binds to 202 sites (Fig. 6d). These data demonstrate that a miRNA can robustly interact with hundreds to thousands of target sites in 3’UTRs even in one cell type. Interestingly, for each of these top 40 miRNAs, we observed 35.5% to 67.2% of miRNA-associated mRNAs contain the seed match found by HOMER. Because seedless sites have a minimal impact on gene expression for both miR-200s and miR-205 (Figs. 2e and 2g), these data may indicate an activity of target scanning by miRNAs that are captured by CLEAR-CLIP.

miRNA-mediated gene expression regulation is highly complex. To date, it remains unclear how many genes and pathways are regulated by miRNAs in a specific cellular context. To gain insights into the overall function of the miRNA pathway in epithelial cells of the skin, we analyzed the targets of these well-expressed miRNAs to determine if different miRNAs work in concert to regulate similar genes or pathways and which genes and signaling pathways are most heavily affected by the miRNA pathway. To accomplish this, we used the same miRNAs with 50 or more high confidence sites as above, calculated high confidence sites within 3’UTRs with a seed match (7mer or better) and determined gene lists for each miRNA. We then performed hierarchical clustering on the table of miRNAs and their targets. As shown in figure S4a, miRNAs that clustered together by genes targeted mostly shared identical seeds, indicating that the identical seed match is the strongest driver for miRNA target coordination. Next, we used Enrichr [25] to classify individual genes into GO terms by KEGG pathways and analyzed whether different miRNAs may regulate similar cellular functions. We found that miRNA families tend to cluster together such as a cluster of let-7 miRNAs (Fig. 6e). However, we observed miR-30b-5p also clustered together with the let-7 family. Additionally, miR-15a/b, miR-16 and miR-29a/b also clustered together. These miRNAs have only one base pair difference in their seed regions and indeed appear to be regulating similar pathways. Interestingly, the miR-200 family, miR-19a, miR-203, miR-301 and miR-27a/b also form a cluster, raising the possibility that different miRNAs coordinately target similar pathways.

To better determine what pathways are strongly targeted by the miRNA pathway globally, we calculated how many miRNAs were targeting each GO term. This analysis revealed that the majority of the top categories targeted by miRNAs in the epithelial cells of the skin are important regulators of cancer (Fig. 6f). In addition, PI3K-Akt signaling, focal adhesion, Hippo pathway and p53 pathways are also strongly targeted.

Because the number of CLEAR-CLIP reads reflects the strength of repression (Fig. 4), we next identified strongly regulated targets by all miRNAs. We calculated the total number of unique miRNAs targeting events for each gene and the total number of CLEAR-CLIP reads per gene. To identify highly targeted genes, we selected for genes that were targeted by 4+ different miRNAs and harbored 40 or more total CLEAR-CLIP reads. To test the effectiveness of this approach, we utilized data from mouse ESCs where endogenous Ago1-4 are deleted and supplemented with an inducible Ago2, allowing acute activation of the entire miRNA pathway [30]. We identified commonly expressed genes (basemean>10) in both systems and determined the expression changes of highly targeted genes upon the activation of the miRNA pathway. Genes with 4+ miRNAs and 40 or more total miRNA reads were more heavily repressed upon activation of the miRNA pathway than all genes targeted by miRNAs, and both were repressed relative to non-targeted genes (Fig. S4b). These data validate our selection method and suggest that genes heavily targeted by the miRNA pathway are similar between cell types when both mRNAs and miRNAs are expressed.

Finally, we performed GO analysis for KEGG terms on genes heavily targeted by the miRNA pathway (Fig. 6g and Table S2). Many of the top pathways were again related to tumorigenesis. The top category was Proteoglycans in Cancer, with 52 out of 203 genes in this category heavily targeted by the miRNA pathway. Three of the top seven categories were also related to cell adhesion including focal adhesion, adherens junction, and regulation of actin cytoskeleton. Taken together, these data offer new insights about miRNA-mediated regulation in epithelial cells of the skin and provide a molecular basis to explore the miRNA pathway as an important negative regulator of tumorigenesis.

## Discussion

In this study, we have optimized CLEAR-CLIP for capturing miRNA:RNA interactions and demonstrated the application of this method for probing the action of miRNAs in epithelial cells at a genomic scale. Although the ligation between miRNA and RNA fragments remains a relatively rare event in comparison to miRNA only and mRNA only events (Table S1), the use of PEG-8000, randomized 3’ and 5’ adapters and enhanced RNA isolation from membrane (Fig. 1a) improves the quantitative performance of CLEAR-CLIP. In support of the notion that more miRNA binding correlates with more robust regulation, our analysis demonstrates that the number of CLEAR-CLIP reads (Figs. 4a and 4d), the percentage of individual miRNA CLEAR-CLIP reads among the total CLEAR-CLIP reads on the same gene (Figs. 4b-c and 4e-f) and the number of unique CLEAR-CLIP identified miRNA binding sites (Fig. S3a-c) all contribute to the strength of miRNA-mediated regulation. These global analyses are also supported by our recent study that individual sites harboring more CLEAR-CLIP reads generally confer stronger regulation as assayed by the classic luciferase assay [18]. The quantitative performance of our CLEAR-CLIP is further validated by the detection of more CLEAR-CLIP reads on the same sites in miR-200 induced epithelial cells than control cells (Figs. 2a and 2c). Importantly, genes commonly targeted in control and miR-200 induced cells are more strongly downregulated than genes uniquely bound by miR-200s in the induced cells (Fig. 2b). These data suggest that elevated miRNA expression preferentially regulates existing targets than *de novo* targets, at least for robustly expressed miRNAs such as miR-200s in epithelial cells.

Direct and quantitative capture of miRNA targets also provides new insights into how miRNAs recognize their targets. Although perfect seed matches, in particular 7mer and 8mer matches, result in the strongest regulation (Fig. 2d-h), miRNAs also bind to a large number of sites that lack a seed match (Figs. 1h and 6d). While most of these sites do not confer discernible regulation to host genes, these observations suggest that miRNAs and their associated RISC may scan a large number of sites within and outside of 3’UTRs, perhaps through an AGO2 phosphorylation-dependent mechanism that has been recently demonstrated [31], but form a more stable complex on high-quality 3’UTR sites. Although miRNAs predominantly use their 5’ seed regions to recognize their targets, base-pairing at the 3’ regions of individual miRNAs has also been described [2,11,13,32]. Analysis of top miRNAs in epithelial cells reveals that different miRNAs have different preferences to the matches in their 3’ regions (Fig. 6b-c). For example, miR-200 miRNAs have two short 3’ regions that help target recognition. In particular, nucleotides 12-14, which are identical in all family members (GGU), show a strong preference to base pair with their targets, providing an explanation for the overlapping targets of different miR-200s with identical seed region (Fig. 3c). In contrast, miR-205 prefers 3’ end binding of nucleotides 15-20 (Fig. 3d). The comparison between CLEAR-CLIP identified and TargetScan predicted targets offers further validation to the experimental approach. In particular, the ability to reduce many false positive predications should help to detect highly regulated targets and their relevant pathways.

Genome-wide identification of miRNA-associated mRNA sites also provide comprehensive understanding of miRNA-controlled transcriptome in a cellular context-specific manner. In the epithelial cells, we have identified >9,000 sites that harbor at least a 7mer or 8mer match to miRNA seed regions (Fig. 5j). Analyses of these targeted genes reveals a picture where epithelial miRNAs regulate numerous genes involved in cancer, focal adhesion, adherens junction, FoxO signaling and Hippo signaling among others (Fig. 6f-g). This knowledge is consistent with genetic studies of *Dicer1, Dgcr8* and *Ago1/2*, in which deletion of these essential factors of the miRNA pathway in epithelial cells of the skin does not change the cell fate but leads to defects in hair morphogenesis and stem cell maintenance [8,33,34]. Although the detailed mechanisms remain to be elucidated, the comprehensive mapping of miRNA targeted transcriptome will lay a foundation to answer the question of how miRNAs regulate hair morphogenesis. These results also provide support to the published reports indicating that miRNA dysregulation is causal in many types of cancer [35–37].

## Conclusion

In conclusion, we have demonstrated the global binding landscape of miRNAs in a miRNA- and site-specific manner and revealed the impact of miRNAs on the transcriptome. Because of the complexity of individual miRNAs’ sequences and their numerous target sites, experimental identification of miRNA targets through optimized CLEAR-CLIP should be the most effective method to detect binding sites for each miRNA. Such holistic analyses of miRNA regulated genes will help to reveal the functions of these small noncoding RNAs in any cellular contexts.

## Methods

### CLEAR-CLIP

Mouse keratinocytes of the designated genotype were maintained in E-low calcium medium. Inducible cells were treated with 3 ug/ml final concentration doxycycline for 24 hours before performing CLEAR-CLIP. One 15cm dish of confluent cells was used per sample. Cells were washed once with cold PBS. 10mls of cold PBS was then added and cells were irradiated with 300mJ/cm^2^ UVC (254nM wavelength). Cells were then scraped from the plates in cold PBS and pelleted by centrifugation at 1,000g for 2 minutes. Pellets were frozen at −80°C until needed. Cells were then lysed on ice with occasional vortexing in 1ml of lysis buffer (50mM Tris-HCl pH 7.4, 100mM NaCl, 1mM MgCl2, 0.1 mM CaCl2, 1% NP-40, 0.5% Sodium Deoxycholate, 0.1% SDS) containing 1X protease inhibitors (Roche #88665) and RNaseOUT (Invitrogen) at 4ul/ml final concentration. Next, TurboDNase (10U), RNase A (0.13ug) and RNase T1 (0.13U) were added and samples were incubated at 37°C for 5 minutes with occasional mixing. Samples were immediately placed on ice and then centrifuged at 16,160g at 4°C for 20 minutes to clear lysate. 25ul of Protein-G Dynabeads were used per IP. Dynabeads were pre-washed with lysis buffer and pre-incubated with 3ul of Wako Anti-Mouse-Ago2 (2D4) antibody. The dynabead/antibody mixture was added to the lysate and rocked for 2 hours at 4°C. All steps after the IP were done on bead until samples were loaded into the polyacrylamide gel. Beads were captured on a magnetic stand and the supernatant removed, then washed 3 times with cold High Salt Clip Wash Buffer (50mM Tris-HCl pH 7.4, 1M NaCl, 1mM EDTA, 1% NP-40, 0.5% sodium deoxycholate, 0.1% SDS) for 3 minutes with rocking. Samples were then washed 2 times with PNK wash buffer (20mM Tris-HCl pH 7.4, 10mM MgCl2, 0.2% Tween-20). Samples were then phosphorylated at 37°C for 20 minutes in 50ul of PNK mixture: 41.8ul H2O, 5 ul 10X PNK buffer (NEB), 1 ul RNaseOUT, 1.67 ul ATP (30 mM), 0.5 ul T4 PNK - 3’ phosphatase minus (NEB M0236L). Samples were then washed 3 times on a magnetic rack with PNK wash buffer. miRNA-mRNA ligation was then carried out overnight at room temperature in 100ul of mixture: 49.25 ul H2O, 30 ul 50% PEG-8000, 10 ul 10X T4 RNA ligation buffer (NEB), 2.5 ul RNaseOUT, 1 ul ATP (100mM), 1 ul BSA (10 mg/ml), 6.25 ul T4 RNA ligase 1 (10U/ul – NEB M0204). Next morning, an additional 2.5 ul T4 RNA ligase 1 (10U/ul) and 1 ul ATP (100mM) were added and ligation was continued for another 5 hours. Samples were then washed 2 times with lysis buffer, once with PNK/EDTA/EGTA (50 mM Tris pH 7.4, 10 mM EDTA, 10 mM EGTA, 0.5% Igepal) and twice more with PNK wash buffer. Next, samples were treated with phosphatase at 37°C for 20 minutes with 50ul of mix: 41ul H2O, 5ul 10X FastAP buffer, 3ul FastAP enzyme (Thermo Fisher #EF0651) and 1ul RNaseOUT. Samples were then washed two times with PNK wash buffer. Next, 3’ adapter ligation was performed on beads overnight in 40 ul of mixture: 17ul H2O, 4ul 10X T4 RNA ligase buffer (NEB), 1ul of 3’linker (5’-Adenylated & 3’ blocked - custom ordered from IDT), 16ul 50% PEG-8000, 1 ul RNaseOUT and 1ul T4 RNA ligase 2 truncated K227Q (NEB M0351). Samples were then washed twice with PNK buffer. Next, samples were then radiolabeled on bead with 50ul of the following mix: 5 ul 10X PNK buffer (NEB), 1 ul RNaseOUT, 1.5 ul y-p^32^-ATP (15μCi), 1 ul PNK enzyme (NEB M0201) and 41.5 ul H2O. Radiolabeling was carried out for 5 minutes at 37°C, after which another 2 ul of cold ATP (10mM) was added and the samples were then incubated for another 5 minutes at 37°C. Samples were then washed three times with PNK buffer, and then re-suspended in 25 ul of 1.2X LDS NuPAGE Loading buffer (Thermo Fisher #NP0007) with 60 mM DTT added. Samples were then heated to 70°C for 10 minutes with occasional agitation and supernatant was separated from beads on a magnetic stand. Samples were loaded on an 8% Bis-Tris gel and run at 200V for 2 hours on ice. Protein-RNA complexes were then transferred to nitrocellulose at 90V for 90min. The membrane was washed with PBS and exposed to a phosphoscreen for 1 hour at −20°C. Fragments corresponding to the Argonaute complex with the miRNA & mRNA (∼110 kDa to 160 kDa) were then excised and RNA was isolated by the following method: 15 ul of Proteinase K at 20 mg/ml was added to 285 ul of Proteinase K/SDS buffer (100 mM Tris-HCl pH 7.5, 50 mM NaCl, 1 mM EDTA, 0.2% SDS). This solution was heated to 37°C for 20 minutes to inactivate any RNases, added to nitrocellulose membrane fragments and incubated at 50°C for 1 hour. Samples were briefly centrifuged and then 375 ul of saturated phenol/choloform/isoamyl alcohol (25:24:1) was added and incubated for 10 minutes at 37°C. Samples were then centrifuged at 16,000g at room temperature for 3 minutes, the aqueous layer was removed to a new tube and precipitated overnight at −20°C with 2 ul Glycoblue (ThermoFisher #AM9516) and 900 ul of 100% ethanol. RNA was pelleted by centrifugation at 16,160g at 4°C for 20 minutes and the supernatant removed. Pellet was washed with 70% ethanol and left to air dry at room temperature for 5 minutes. Next, 5’ adapters were ligated by adding 8ul of the following mix to the pellet and thoroughly resuspending: 2 ul 50% PEG-8000, 1ul 10X NEB RNA ligation buffer, 1 ul 10mM ATP, 4 ul H2O. Samples were then heated briefly to 95°C, placed on ice and additional components were added: 0.5 ul RNaseOUT, 1 ul T4 RNA ligase 1 (10U/ul – NEB M0204), 0.5 ul 100 uM 5’ RNA linker (Blocked at 5’ end & contains NNNN at 3’ end for barcoding). Ligation was carried out at 37°C for 4 hours with rocking. RT-PCR was then carried out in ligation buffer by adding 8.5 ul of the following mix to the sample: 4 ul 5X first-strand buffer (Invitrogen), 1.5 ul 100mM DTT, 2ul 1 uM RT primer, 1 ul 10mM dNTPs. Samples were heated to 65°C for 5 minutes, transferred to a PCR tube and then enzymes were added: 1 ul Superscript III (Thermo Fisher #18080093) and 0.5 ul RNaseOUT. RT reaction was then performed in a thermocycler: 50°C for 1 hour, 85°C for 5 minutes, and then hold at 4°C. Libraries were then amplified from the cDNA by PCR taking aliquots after cycles 12, 17 and 22. PCR mix: 18.8 ul H2O, 8 ul 5X HF buffer (NEB), 1 ul 25 uM library 1^st^ round forward primer, 1 ul 25 uM RT primer, 0.8 ul 10 mM dNTPs, 0.4 ul Phusion polymerase (NEB M0530) and 10 ul cDNA. Cycling parameters: Initial denaturation at 98°C for 30 seconds and then amplification cycles at 98°C for 15 seconds, 56°C for 30 seconds and then 72°C for 20 seconds. PCR products were run on a 9% acrylamide gel and then stained with SYBR gold (ThermoFisher #211494) at 1:10,000 for 10 minutes. The area corresponding to approximately 73 to 150 base pairs was excised from the lowest cycles condition that showed a product. Gel pieces were then frozen at −80°C for one hour and centrifuged through a hole in the tube made with a 20G needle to break up the gel. 400 ul HSCB buffer (25 mM Tris-HCl pH 7.5, 400 mM NaCl, 0.1% SDS) was added and samples were then rocked overnight at 4°C. The next day the gel slurry was transferred to a 0.22 um filter tube and spun at 16,000g for 20 minutes at room temperature. Samples were then precipitated overnight at −20°C in the presence of 1 ml 100% ethanol and 2ul Glycoblue. The next day samples were centrifuged at 16,160g at 4°C for 20 minutes. The pellet was washed with 70% ethanol and then air dried for 5 minutes. The pellet was then resuspended in 20 ul of H2O. Next, high throughput sequencing barcodes were added by PCR using the following mix: 10 ul previous PCR product, 3.84 ul H2O, 4 ul 5X HF buffer, 0.66 ul 10mM dNTPs, 0.5 ul 25 uM Illumina Index Primer, 0.5 ul 25 uM Illumina RP1 primer and 0.5 ul Phusion. Cycling conditions: Initial Denaturation at 98°C for 30 seconds, 2 cycles of: 98°C for 15 seconds, 50°C for 20 seconds, 72°C for 45 seconds and then 4 cycles of: 98°C for 15 seconds and 72°C for 50 seconds and then a final extension at 72°C for 3 minutes. PCR products were then run on a 9% acrylamide gel and stained with SYBR gold as above. Products corresponding to sizes approximately 144-200 base pairs were excised and isolated from the gel as above. Libraries were mixed in equal amounts and sequenced on an Illumina HiSeq 4000 by the Microarray and Genomics Core at the University of Colorado Anschutz Medical Campus. Primers and adapters used for CLEAR-CLIP were published previously[18].

### CLEAR-CLIP samples

More details given in Supplemental Table 1. CLEAR-CLIP was done on 18 libraries in total from mouse keratinocytes. 9 control libraries (6 K14-Cre only cells, and 3 miR-200 inducible cells not treated with doxycycline), 6 miR-200 DKO cells, and 3 miR-200 inducible cells (treated for 24 hours with 3 ug/ml Doxycycline final concentration.

### Assigning Chimeric Reads and Genome Annotation for CLEAR-CLIP

This bioinformatic analysis was done the same as previously[18].

### Cells

Cells were isolated and maintained as previously described[18].

### Defining high confidence areas

High confidence areas were defined using mRNA areas specific to each miRNA. Areas for each strand were separated and run through MultiIntersectBed, as the program is not strand aware. Areas found in 2+ libraries were then selected and merged together. Areas were then overlapped with 3’UTRs using Bedtools intersect and a database of mm10 3’UTRs downloaded from the USCS table browser. If areas were to be used for motif finding and they were less than 60bp, an equal number of bases was added in each direction to make them 61 or 62 base pairs. Data using 2+, 5+ and 8+ libraries were done similarly, but using the indicated number of libraries. Annotations to the genome were done using Bedtools intersect and genomic annotation sets downloaded from the USCS table browser. In the case of an area overlapping multiple annotations the area was annotated by the following preference: (miRNA > 3’UTR > CDS > 5’UTR > Intron > tRNA > ncRNA). For motif calling genes were assigned by the best miRNA motif they contained (8mer > 7merM8 > 7merA1 > 6mer).

### Unbiased motif finding using Homer

Homer was downloaded from http://homer.ucsd.edu/homer/motif/. High confidence areas found in 2+ libraries were calculated for the top 40 miRNAs, and selected for areas overlapping 3’UTRs using Bedtools. All 3’UTR areas for each miRNA were then processed through Homer. The background for motif finding was all mouse mm10 3’UTRs. Settings for motif finding were: -len 8 -size given -rna -noweight - p 2 -chopify. The top ranked motif is shown for each miRNA.

### RNAhybrid

RNAhybrid was downloaded from https://bibiserv.cebitec.uni-bielefeld.de/rnahybrid?id=rnahybrid_view_download. RNAhybrid was run on areas of interest using using a fasta file of the miRNA sequence. For most experiments it was run using the options -b 1 -s 3utr_human. For the k-means clustering of miR-200s, high confidence areas were split into seed containing regions versus non-seed containing and the -f 2-7 option was used in addition to require seed use. Python scripts were used to parse the RNAhybrid output for either binding energy or binding fractions. Outputs of RNAhybrid were combined into a table of each site with 1 indicating bound or 0 indicating not bound. Data was then k-means clustered using Cluster (https://www.encodeproject.org/software/cluster/). We examined using 4-15 clusters, but we found 5 to be the most informative. Clustered data was then drawn using Java Tree View. Only areas with a motif were used if a gene had both motif containing and non-motif containing areas. Due to the difficulty of determining which group is better *a priori*, if a gene had multiple motif containing areas it could be contained in multiple groups.

### GO terms clustering

GO terms were calculated using Enrichr https://amp.pharm.mssm.edu/Enrichr/. The KEGG 2016 categories were used for GO Terms and the combined score was used as output for graphing. Scores were then hierarchal clustered using Gene Cluster 3.0 (https://www.encodeproject.org/software/cluster/) and drawn as a heat map using Java Tree View (http://jtreeview.sourceforge.net/).

### qPCR

miRNA qPCR was performed using the Qiagen miScript II RT Kit (218160). miRNAs were quantified from the cDNA using iQ SYBR green supermix (170-8880; Bio-Rad Laboratories) and the ΔΔC(t) method relative to U6 RNA. Forward primers for miRNAs are as follows: miR-205 5’ TCCTTCATTCCACCGGAGTCTG 3’, miR-200a 5’ TAACACTGTCTGGTAACGATGT 3’, miR-200b 5’ TAATACTGCCTGGTAATGATGA 3’, miR-200c 5’ TAATACTGCCGGGTAATGATGGA 3’, miR-141 5’ TAACACTGTCTGGTAAAGATGG 3’, miR-429 5’ TAATACTGTCTGGTAATGCCGT 3’. Reverse primer used was Qiagen’s miScript Universal Primer.

### RNA-seq

RNA-seq was performed on both miR-200b cluster induced cells and miR-205 induced cells. Controls for both were matching cells, but not induced with Doxycycline, just treated with vehicle. Doxycycline treatment was 3ug/ml for 24 hours for both cells. RNA was harvested in trizol from 6 well plate for each sample. miR-200 induction RNA-seq was performed in duplicate. miR-205 induction RNA-seq was performed in triplicate. RNA was isolated from Trizol using the standard method. RNA was then poly(A) selected using the ambion Dynabead mRNA DIRECT Purification kit (#61012). Library preparation was performed using the NEBNext Ultra Directional RNA Library Prep Kit for Illumina (#E7420). Sequencing reads were mapped to the mm10 genome using Bowtie2 and gene counting performed using HTSeq-count. Differential analysis was performed using DESeq2.

### Non-coding RNA studies

Non-coding RNAs were downloaded from noncode.org. This database has both “genes” and “transcripts,” herein the set of “genes” was used. RNA-seq data was intersected with the database of non-coding RNAs using HTSeq-count. DESeq2 was then used to calculate fold change data. For motif calling genes were assigned by the best miRNA motif they contained (8mer > 7merM8 > 7merA1 > 6mer). For comparing location of the sites, ncRNAs that were down more than 0.1 Log2 fold were deemed “effective” and all other ncRNAs were deemed “ineffective.”

### TargetScan Comparisons

All comparisons were done to TargetScan Mouse 7.1, using their database of conserved sites. These were compared to CLEAR-CLIP data for genes with a high confidence site that contained an 8mer or either 7mer (M8 or A1). Comparisons were done by gene, irrespective of whether the identical site was found in both. Comparisons were also only done for genes that had a basemean of >10 in our RNA-seq data to make sure we compared genes that were expressed in our system.

### Statistics

All measurements were taken from separate samples. All statistical tests used are described in the figure legends. For graphing of Log2 data, such as figure 2a, 1 was added to each data point to make the Log2 of 0 possible.

### Code Availability

Most analyses were performed using publicly available programs such as Bedtools. Custom scripts such as CLEAR-CLIP mapping steps, Bedfile area extending, Blast output processing and scripts to parse RNAhybrid are available at https://github.com/Bjerkega/CLEAR-CLIP-analysis-scripts.

## Declarations

### Availability of data and material

High throughput sequencing data is available online from the Gene Expression Omnibus (GEO) at GSE102716 (CLEAR-CLIP) and GSE131205 (RNA-seq).

### Authors’ contributions

G.A.B. and R.Y. conceived of the study and wrote the manuscript. G.A.B. performed the experiments and analyses.

## Acknowledgments

We thank K. Diener and B. Gao for sequencing; J. Hoefert, D. Wang and J. Wang for generating miR-200 and miR-205 mouse models and isolating primary cells; and members of the Yi laboratory for discussion.

