## Supplemental Figures and Tables for "Integrated analysis of directly captured microRNA targets reveals the impact of microRNAs on mammalian transcriptome"

#### Supplemental Figure Legend

**Figure S1. Further characterization of CLEAR-CLIP results. a**, Sequence diagram of murine members of the miR-200 family showing the two related seed sequences and conservation in the 3' end. **b**, qPCR validation of RNA used for RNA-seq showing miRNA levels after miR-200b cluster induction. Shown as the average of biological triplicates +/- the standard deviation. **c**, qPCR validation of RNA used for RNA-seq showing miR-205 level after miR-205 induction. Shown as the average of biological triplicates +/- the standard deviation. **d**, Log2 fold change in gene expression following miR-200b cluster induction of genes found only in CLEAR-CLIP, only in HITS-CLIP or found in both. Only CLEAR-CLIP genes with a seed (6mer or better) were used since HITS-CLIP requires a seed for determining targets. **e**, Log2 fold change in gene expression upon miR-205 induction of genes found only in CLEAR-CLIP (with a seed), only in HITS-CLIP or found in both. **f**, Log2 fold change in gene expression following miR-200b cluster induction of genes that had a CLEAR-CLIP area in only one gene area (3'UTR, 5'UTR, CDS or Intron). Panels **d, e & f:** Number of genes are shown in parenthesis for each group and p-values were calculated using the Kolmogorov–Smirnov test. **g**, Percent overlap of miR-200 members within individual areas is shown and whether the overlap was within or across seed type families. **h**, Full table of miR-200 GO term clustering is shown, including GO categories that were only observed for one miR-200 family member.

**Figure S2. Characterization of CLEAR-CLIP by gene expression level and effect on non-coding RNAs. a**, Log2 gene expression levels are shown for genes with CLEAR-CLIP reads from miRNAs other than miR-200s that were found to be derepressed or not upon induction of the miR-200b cluster. **b**, The log2 number of CLEAR-CLIP reads is shown for genes with CLEAR-CLIP reads from miRNAs other than miR-200s that were found to be derepressed or not upon induction of the miR-200b cluster. P-values for panels **a & b** were calculated using an unpaired two-sided t-test. **c**, The top motif found by HOMER is shown for miR-205 CLEAR-CLIP areas that did not have a 6mer or better canonical motif. **d**, Examples of non-coding RNAs that were found to be targeted by miR-200s are shown for control, miR-200 DKO and miR-200 Tg samples. Fold change data upon induction of the miR-200b cluster is also shown on the right for important non-coding RNAs. **e**, The distance from each miR-200 CLEAR-CLIP site to the 3' end of non-coding RNAs was calculated and shown here as a histogram for non-coding RNAs that were found to be repressed (>0.1 Log2 fold change) upon miR-200b cluster induction (effective) or not (ineffective). The x axis has been clipped for clarity.

**Figure S3. Gene repression by the number of CLEAR-CLIP areas per gene and gene expression of areas found by CLEAR-CLIP or TargetScan. a**, Log2 fold change in gene expression levels upon induction of the miR-200b cluster are shown for genes with 1, 2, 3 or 4+ miR-200 CLEAR-CLIP areas. **b**, Log2 fold change in gene expression levels upon induction of the miR-200b cluster are shown for genes with 1, 2 or 3+ miR-200 CLEAR-CLIP areas with a miR-200 seed. **c**, Log2 fold change in gene expression levels upon induction of miR-205 are shown for genes with 1, 2 or 3+ miR-205 CLEAR-CLIP areas. Panels **a, b & c**: The number of genes in each group are shown in parenthesis and p-values were calculated using the Kolmogorov–Smirnov test. **d**, Log2 gene expression levels as measured by RNA-seq are shown for miR-200 and miR-205 targets that were found only in CLEAR-CLIP, only by TargetScan or found in both.

**Figure S4. Global miRNA targeting analysis. a**, Hierarchical clustering of top miRNAs based on their CLEAR-CLIP gene targets. **b**, Highly targeted genes were defined in our CLEAR-CLIP for genes that were targeted by 4 or more miRNAs and also had 40 or more CLEAR-CLIP reads per gene. Shown here is log2 fold change in gene expression for highly targeted genes upon induction of Ago2 in a mESC cells that were otherwise Ago1-4 null (requiring the genes to be expressed in our mouse keratinocyte RNAseq and their mESC RNAseq). The number of genes for each group is shown in parenthesis and p-values were calculated using the Kolmogorov–Smirnov test.

**Table S1.** List of samples used for CLEAR-CLIP. Table contains the sample type, experiment, the number of sequencing reads and number of CLEAR-CLIP reads.

**Table S2.** Full table showing GO term results for miRNAs that are heavily targeted. The table shows the GO term, the overlap, significance and the genes within the GO term.

#### Supplemental figure 1. Further characterization of CLEAR-CLIP results

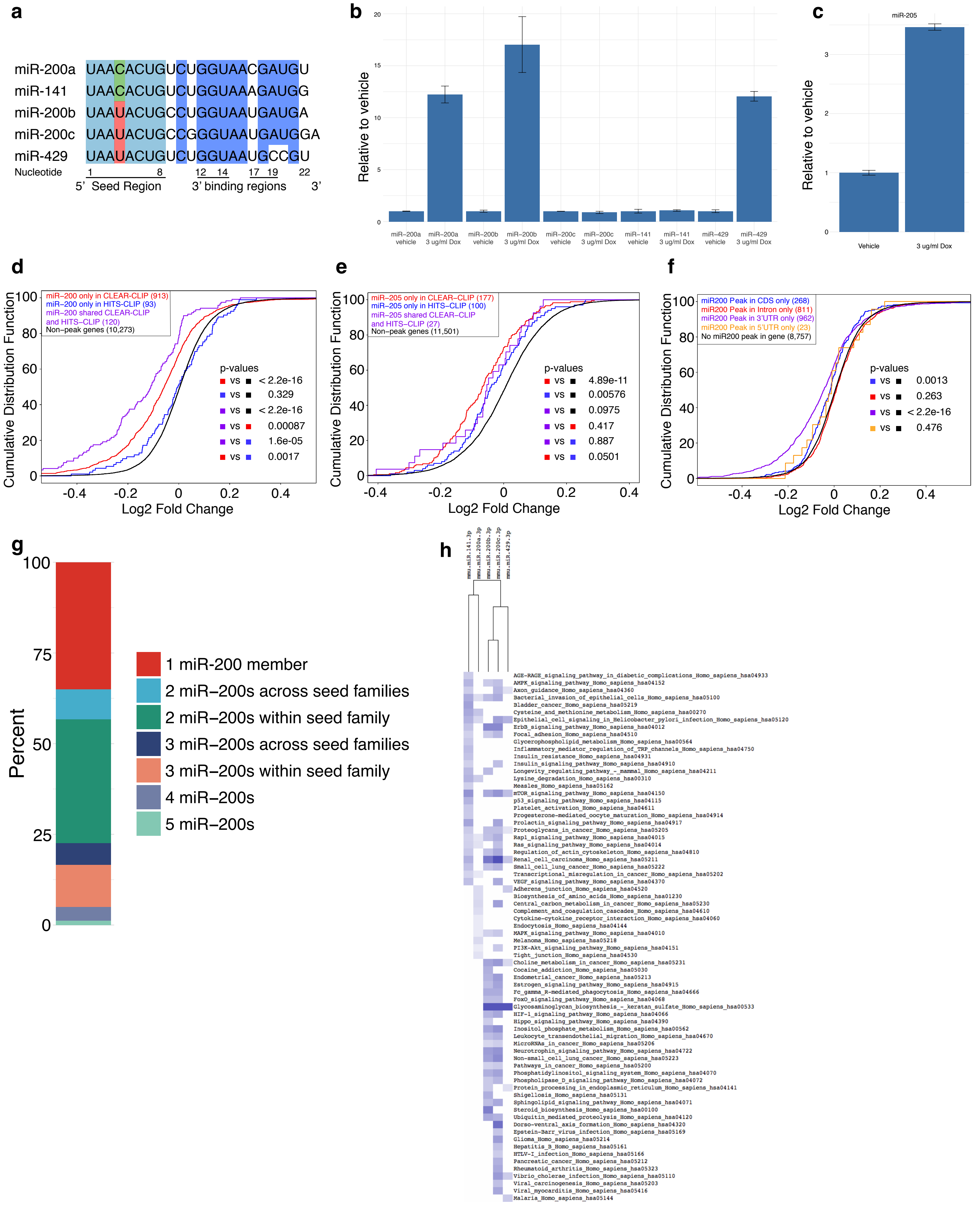

**Supplemental figure 2. Characterization of CLEAR-CLIP by gene expression and effect on ncRNAs**

**a**

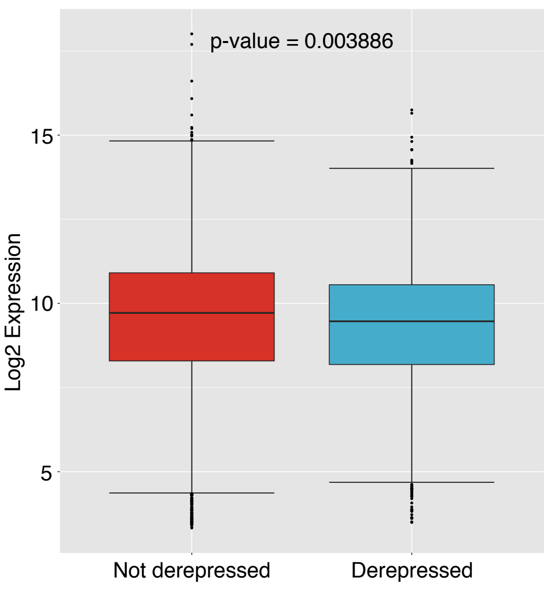

**b**

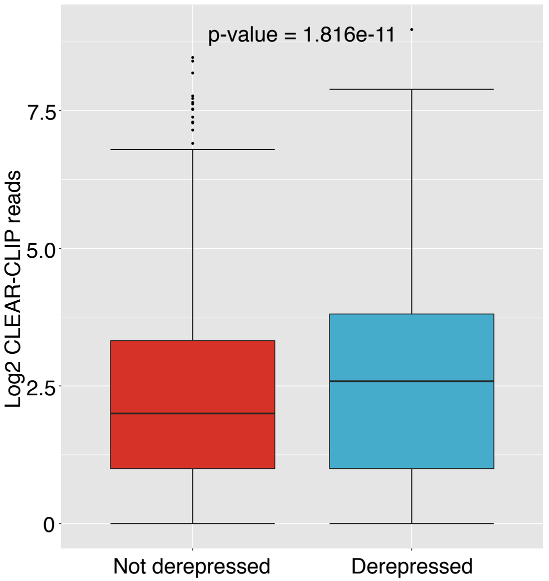

**c**

| Motif | P-value | log P-pvalue | % of Targets | % of Background |
|---|---|---|---|---|
| 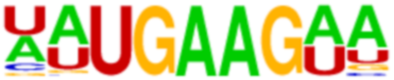 | 1e-415 | -9.557e+02 | 21.79% | 3.45% |

**d**

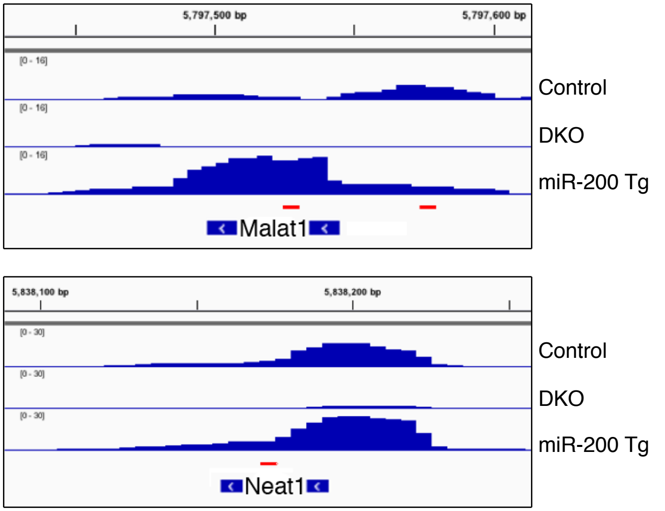

| Log2 Fold Change | |
|---|---|
| Malat1 | -0.24 |
| Neat1 | -0.12 |
| Tug1 | -0.04 |
| Gas5 | -0.05 |
| Firre | -0.03 |

**e**

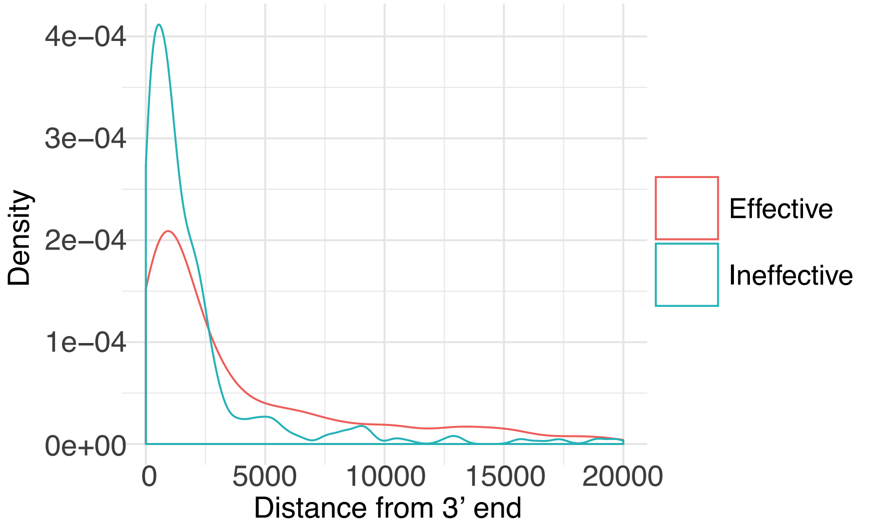

**Supplemental figure 3. Gene repression by the number of CLEAR-CLIP areas per gene and expression levels of genes found by CLEAR-CLIP or TargetScan**

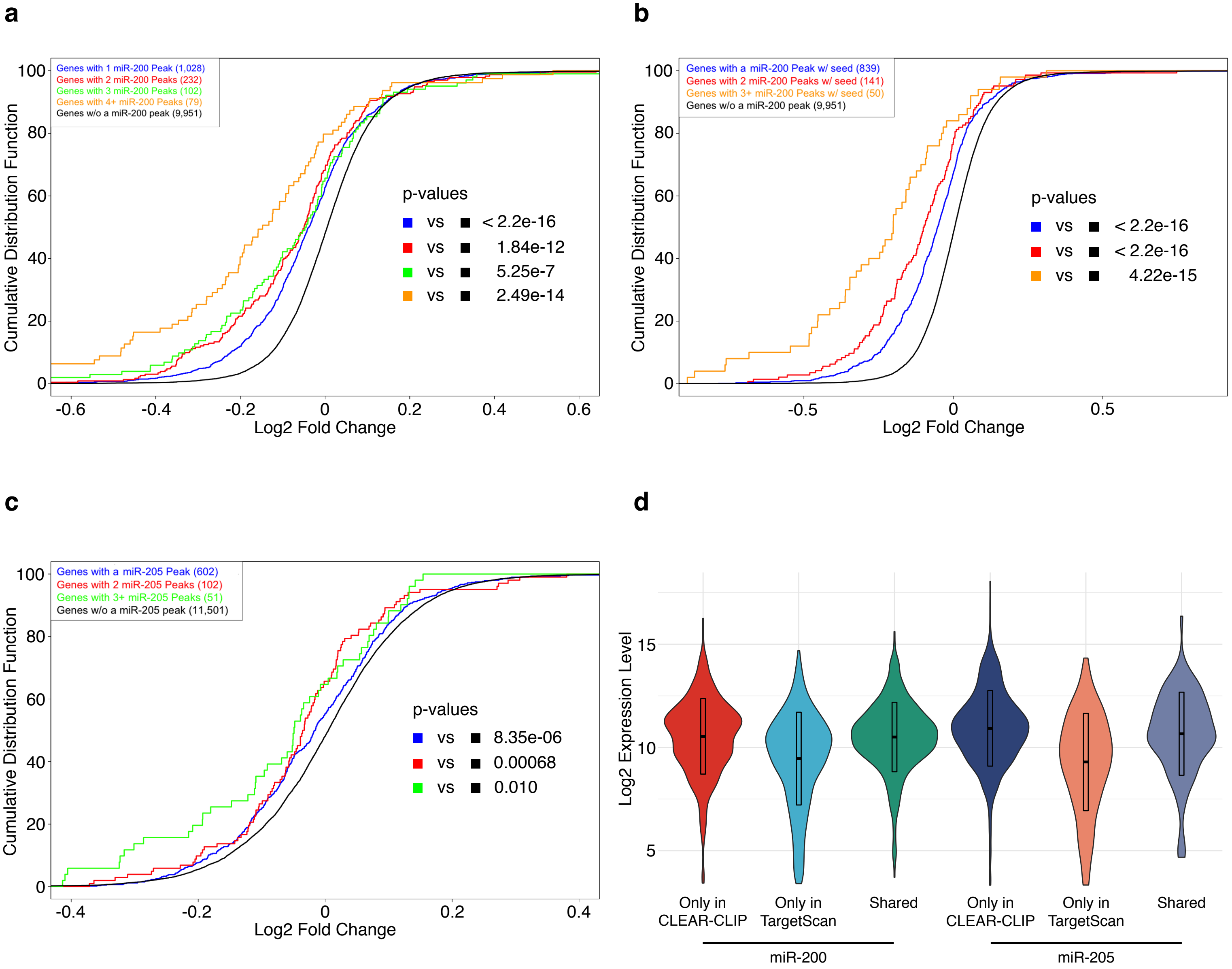

#### Supplemental figure 4. Global miRNA targeting analysis

**a**

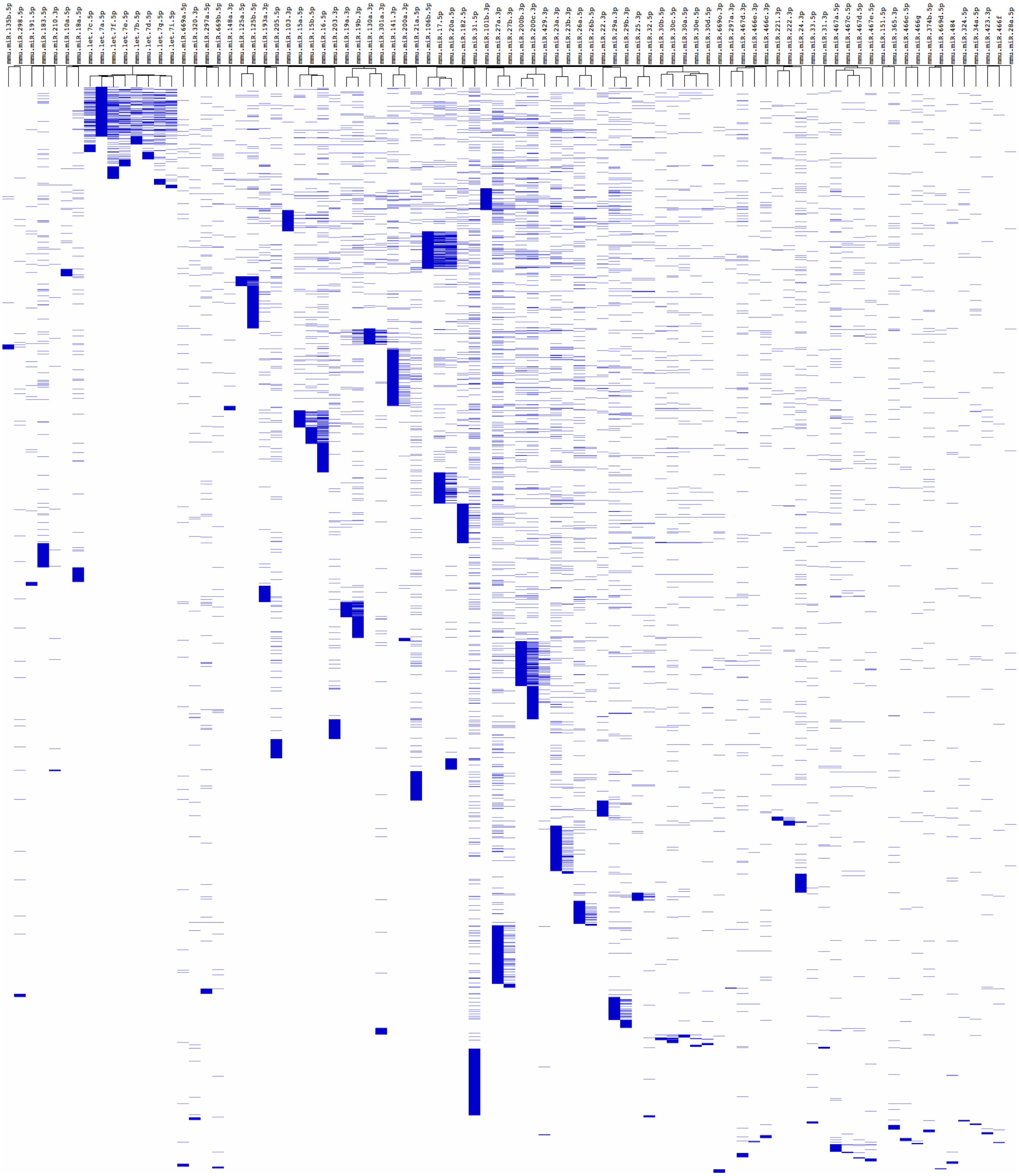

**b**

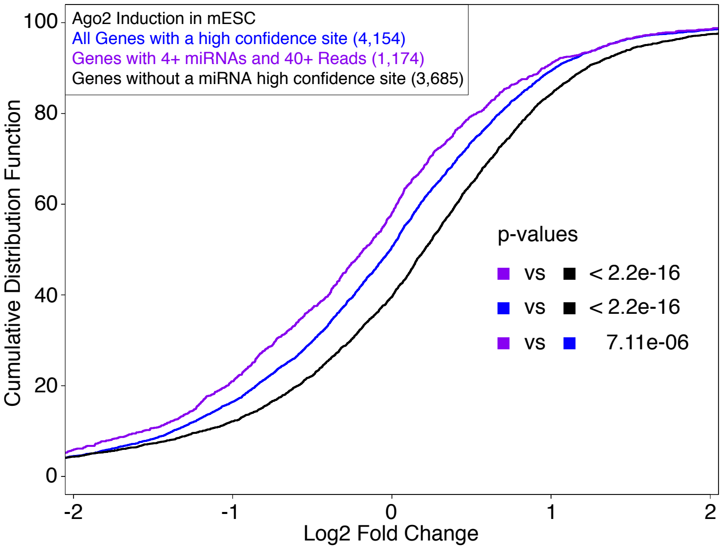

#### Supplementary Table 1. Samples used for CLEAR-CLIP

| Sample | Type | Experiment | Total sequencing reads | Number of unique CLEAR-CLIP reads |
|---|---|---|---|---|
| K14-Cre keratinocytes (Control) | Control 1 | Group 1 | 28,951,235 | 22,394 |
| K14-Cre keratinocytes (Control) | Control 2 | Group 1 | 69,562,643 | 19,308 |
| K14-Cre keratinocytes (Control) | Control 3 | Group 1 | 58,490,875 | 36,400 |
| K14-Cre keratinocytes (Control) | Control 4 | Group 1 | 36,321,840 | 37,567 |
| K14-Cre keratinocytes (Control) | Control 5 | Group 1 | 32,850,327 | 33,594 |
| K14-Cre keratinocytes (Control) | Control 6 | Group 1 | 47,737,230 | 33,483 |
| miR-200 DKO keratinocytes | miR-200 DKO 1 | Group 1 | 36,217,636 | 23,204 |
| miR-200 DKO keratinocytes | miR-200 DKO 2 | Group 1 | 19,825,464 | 16,620 |
| miR-200 DKO keratinocytes | miR-200 DKO 3 | Group 1 | 26,394,791 | 11,596 |
| miR-200b cluster inducible keratinocytes (Un-induced) | Control 1 | Group 2 | 39,414,925 | 57,279 |
| miR-200b cluster inducible keratinocytes (Un-induced) | Control 2 | Group 2 | 41,967,583 | 143,874 |
| miR-200b cluster inducible keratinocytes (Un-induced) | Control 3 | Group 2 | 35,724,788 | 127,034 |
| miR-200b cluster inducible keratinocytes | Inducible 1 | Group 2 | 41,895,809 | 147,001 |
| miR-200b cluster inducible keratinocytes | Inducible 2 | Group 2 | 69,418,670 | 150,488 |
| miR-200b cluster inducible keratinocytes | Inducible 3 | Group 2 | 24,074,912 | 152,936 |
| miR-200 DKO keratinocytes | miR-200 DKO 1 | Group 2 | 51,932,214 | 79,810 |
| miR-200 DKO keratinocytes | miR-200 DKO 2 | Group 2 | 27,092,199 | 103,480 |
| miR-200 DKO keratinocytes | miR-200 DKO 3 | Group 2 | 30,629,688 | 33,951 |
| | | Total | 718,502,829 | 1,230,019 |

### Supplementary Table 2. GO terms associated with miRNA targets

| Term | Overlap | P-value | Adjusted P-value | Old P-value | Old Adjusted P-value | Z-score | Combined Score | Genes |
|---|---|---|---|---|---|---|---|---|
| Proteoglycans in cancer_Homo sapiens_hsa05205 | 54/203 | 2.234E-17 | 5.51799E-15 | 5.16123E-11 | 1.27482E-08 | -2.013336866 | 77.19163974 | ITGB1;CDKN1A;CBLB;PIK3CB;ACTB;IGF1R;PPP1CB;CCND1;CTSL;PLAU;MYC;ITGAV;ARHGEF12;PPP1R12A;PDPK1;GAB1;PLAUR;RRAS2;FRS2;HSPG2;TIAM1;CTTN;PIK3CA;ITGA5;EZR;MET;CD44;HBEGF;DDX5;SDC4;ROCK2;SRC;PIK3R1;IQGAP1;HIF1A;THBS1;EGFR;CDC42;NRAS;MAPK1;PLCG1;SMAD2;CAV2;CAV1;FZD6;RDX;WNT7A;MSN;BRAF;VEGFA;RPS6KB1;MDM2;SDC1;CTNNB1 |
| Focal adhesion_Homo sapiens_hsa04510 | 47/202 | 5.89789E-13 | 4.86156E-11 | 3.62364E-08 | 2.98346E-06 | -1.896862124 | 53.41376404 | ITGB1;GSK3B;ROCK2;SRC;LAMA3;PTEN;XIAP;LAMC2;PIK3R1;PIK3CB;LAMC1;THBS1;ACTB;EGFR;CRKL;IGF1R;RAP1B;PPP1CB;CDC42;CCND3;MAPK8;CCND2;CCND1;CAPN2;PAK6;MAPK1;ITGAV;ITGB6;PAK2;JUN;PPP1R12A;CAV2;ITGA3;PDPK1;ACTN1;CAV1;BRAF;VEGFA;COL4A2;PIK3CA;COL4A1;CTNNB1;ITGA6;ITGA5;CRK;MET;VCL |
| Adherens junction_Homo sapiens_hsa04520 | 27/74 | 5.90472E-13 | 4.86156E-11 | 2.10495E-08 | 2.59962E-06 | -1.803594707 | 50.78535563 | SRC;CTNND1;PTPRJ;IQGAP1;WASL;NLK;MLLT4;PTPRF;EGFR;ACTB;IGF1R;CDC42;CDH1;MAPK1;SMAD2;ACTN1;INSR;TGFBR1;TGFBR2;TJP1;SNAI2;CTNNB1;PVRL4;PVRL2;MET;PVRL1;VCL |
| FoxO signaling pathway_Homo sapiens_hsa04068 | 36/133 | 2.71921E-12 | 1.34329E-10 | 6.57695E-08 | 3.57372E-06 | -1.724599199 | 45.92724875 | CDKN1A;CDKN1B;FOXG1;SETD7;PTEN;PIK3R1;PIK3CB;FOXO3;NLK;STK4;EGFR;IGF1R;NRAS;MAPK8;BCL2L11;CCND2;CCND1;MAPK1;SMAD2;PLK3;HOMER1;CHUK;PDPK1;INSR;PLK2;BRAF;FBXO32;TGFBR1;TGFBR2;PIK3CA;BCL6;CCNG2;CDK2;MDM2;ATM;SGK3 |
| Hippo signaling pathway_Homo sapiens_hsa04390 | 39/153 | 2.6664E-12 | 1.34329E-10 | 7.23425E-08 | 3.57372E-06 | -1.664311085 | 44.3543769 | YAP1;YWHAE;GSK3B;BMPR2;YWHAB;SERPINE1;PPP2R2A;ACTB;CTGF;PPP1CB;PPP2CB;PARD6B;CCND3;CCND2;CCND1;CDH1;MYC;TEAD1;YWHAG;SMAD2;TEAD4;WWTR1;PRKCI;FBXW11;FZD6;WNT7A;CSNK1D;YWHAZ;TGFBR1;SMAD7;TGFBR2;MOB1B;LATS1;DLG1;LATS2;SNAI2;CTNNB1;AJUBA;BMPR1A |
| Regulation of actin cytoskeleton_Homo sapiens_hsa04810 | 46/214 | 1.97943E-11 | 6.98455E-10 | 3.74286E-07 | 1.15561E-05 | -1.772519354 | 43.68485414 | ITGB1;NCKAP1;ROCK2;SRC;ARPC1A;PIK3R1;IQGAP1;WASL;PIK3CB;ACTB;EGFR;CRKL;PPP1CB;CDC42;NRAS;CFL2;PAK6;MAPK1;PIP4K2B;ITGAV;ITGB6;PAK2;ARHGEF12;PPP1R12A;ITGA3;ACTN1;LIMK2;RDX;RRAS2;MSN;BRAF;GNG12;SSH2;ENAH;TIAM1;DIAPH2;PIKFYVE;PIK3CA;MYH9;ITGA6;ITGA5;PFN1;EZR;CRK;VCL;FGFR2 |
| Protein processing in endoplasmic reticulum_Homo sapiens_hsa04141 | 40/169 | 1.75587E-11 | 6.98455E-10 | 2.65583E-07 | 9.37173E-06 | -1.694934096 | 41.97583829 | TRAM1;SEC23A;RPN2;PRKCSH;SEL1L;CUL1;DERL1;HSP90B1;UBE2J1;ERO1L;SEC61A2;ATXN3;MAPK8;GANAB;HSPH1;MAN1A2;LMAN2;CAPN2;SSR1;SEC63;PDIA3;EDEM3;HSPA8;SEC24A;HSPA5;EDEM1;SSR2;SSR3;SYVN1;MOGS;YOD1;UBE2G1;MARCH6;PDIA6;CKAP4;DNAJC5;CANX;ERP29;HYOU1;NFE2L2 |
| p53 signaling pathway_Homo sapiens_hsa04115 | 24/69 | 3.69495E-11 | 1.14082E-09 | 2.65595E-07 | 9.37173E-06 | -1.529528901 | 36.74153062 | CDKN1A;RRM2;APAF1;EI24;IGFBP3;SERPINE1;PTEN;TNFRSF10B;PPM1D;THBS1;SERPINB5;CCND3;CDK6;CCND2;CCND1;ZMAT3;CCNG2;PERP;CCNG1;CDK2;MDM2;PMAIP1;ATM;MDM4 |
| Pathways in cancer_Homo sapiens_hsa05200 | 67/397 | 7.07619E-11 | 1.94202E-09 | 2.28425E-06 | 4.86568E-05 | -1.896231922 | 44.31816407 | ITGB1;GSK3B;CDKN1A;CDKN1B;PTEN;SLC2A1;CBLB;LAMC2;PIK3CB;LAMC1;CRKL;IGF1R;CCND1;CDH1;MYC;ITGAV;ARHGEF12;CHUK;ITGA3;FOS;TGFBR1;RUNX1;TGFBR2;COL4A2;PIK3CA;COL4A1;TRAF6;ITGA6;CRK;MET;ROCK2;LAMA3;GNAI3;LPAR2;TGFA;XIAP;PIK3R1;PTGS2;STK4;HIF1A;ADCY6;EGFR;HSP90B1;CDC42;NRAS;MAPK8;E2F2;MAPK1;E2F3;PLCG1;SMAD2;JUN;EGLN3;JUP;FZD6;WNT7A;BRAF;GNG12;VEGFA;CDK6;LPAR6;CDK2;CCDC6;MDM2;GNAS;CTNNB1;FGFR2 |
| Bacterial invasion of epithelial cells_Homo sapiens_hsa05100 | 25/78 | 1.10566E-10 | 2.73098E-09 | 5.33336E-07 | 1.46371E-05 | -1.510833424 | 34.63647268 | ITGB1;SRC;ARPC1A;CLTC;CBLB;PIK3CB;PIK3R1;WASL;ACTB;CRKL;CDC42;CDH1;SEPT11;CAV2;CAV1;GAB1;SEPT2;CD2AP;PIK3CA;CTTN;CTNNB1;ITGA5;CRK;MET;VCL |
| ErbB signaling pathway_Homo sapiens_hsa04012 | 26/87 | 2.68955E-10 | 6.03927E-09 | 9.62207E-07 | 2.37665E-05 | -1.60286839 | 35.32157063 | GSK3B;CDKN1A;CDKN1B;SRC;TGFA;CBLB;PIK3CB;PIK3R1;EGFR;CRKL;NRAS;MAPK8;MYC;PAK6;ABL2;MAPK1;PLCG1;PAK2;JUN;GAB1;BRAF;EREG;PIK3CA;RPS6KB1;CRK;HBEGF |
| PI3K-Akt signaling pathway_Homo sapiens_hsa04151 | 59/341 | 3.59601E-10 | 7.40178E-09 | 4.87208E-06 | 8.0227E-05 | -1.781845463 | 38.74806006 | YWHAE;ITGB1;ATF2;GSK3B;CDKN1A;CDKN1B;YWHAB;PTEN;LAMC2;PPP2R2A;PIK3CB;LAMC1;IGF1R;CCND3;CCND2;CCND1;MYC;CREB3L2;ITGAV;ITGB6;YWHAG;CHUK;ITGA3;PDPK1;YWHAZ;CREB1;COL4A2;PIK3CA;COL4A1;DDIT4;SGK3;ITGA6;ITGA5;MET;EPHA2;IFNAR1;PHLPP2;LAMA3;LPAR2;PIK3R1;EFNA5;FOXO3;THBS1;EGFR;HSP90B1;PPP2CB;NRAS;BCL2L11;MAPK1;MCL1;INSR;GNG12;VEGFA;CDK6;RPS6KB1;LPAR6;CDK2;MDM2;FGFR2 |
| Small cell lung cancer_Homo sapiens_hsa05222 | 25/86 | 1.12994E-09 | 2.14689E-08 | 2.36389E-06 | 4.86568E-05 | -1.51033928 | 31.11465026 | ITGB1;CDKN1B;LAMA3;PTEN;XIAP;LAMC2;PIK3CB;LAMC1;PIK3R1;PTGS2;CCND1;MYC;E2F2;E2F3;ITGAV;APAF1;CHUK;ITGA3;CDK6;PIK3CA;COL4A2;COL4A1;TRAF6;CDK2;ITGA6 |
| Prostate cancer_Homo sapiens_hsa05215 | 25/89 | 2.49022E-09 | 4.28831E-08 | 3.95279E-06 | 7.5103E-05 | -1.590800867 | 31.51519173 | GSK3B;CDKN1A;CDKN1B;PTEN;TGFA;PIK3CB;PIK3R1;EGFR;HSP90B1;IGF1R;NRAS;CCND1;CREB3L2;E2F2;MAPK1;E2F3;CHUK;PDPK1;BRAF;CREB1;PIK3CA;CDK2;MDM2;CTNNB1;FGFR2 |
| MicroRNAs in cancer_Homo sapiens_hsa05206 | 52/297 | 2.60423E-09 | 4.28831E-08 | 1.36979E-05 | 0.000211461 | -1.570451519 | 31.0417446 | CDKN1A;CDKN1B;BMPR2;PTEN;GLS;CRKL;CCND2;CCND1;PLAU;MYC;PIM1;TPM1;DNMT3A;KIF23;DICER1;SERPINB5;CDC25A;CDC25B;FOXP1;PIK3CA;DDIT4;FSCN1;ITGA5;EZR;CRK;MET;CD44;NOTCH2;NOTCH1;PTGS2;SLC7A1;THBS1;EGFR;NRAS;BCL2L11;IGF2BP1;E2F2;HMOX1;MAPK1;E2F3;PLCG1;MCL1;RDX;HMGA2;VEGFA;MARCKS;CDK6;CCNG1;MDM2;SPRY2;ATM;MDM4 |
| Renal cell carcinoma_Homo sapiens_hsa05211 | 21/66 | 3.94345E-09 | 6.0877E-08 | 4.756E-06 | 8.0227E-05 | -1.382324 | 26.74964095 | JUN;EGLN3;GAB1;SLC2A1;TGFA;BRAF;PIK3CB;PIK3R1;HIF1A;VEGFA;CRKL;RAP1B;CDC42;FLCN;NRAS;PIK3CA;PAK6;MAPK1;PAK2;CRK;MET |
| Endocytosis_Homo sapiens_hsa04144 | 46/259 | 1.35428E-08 | 1.96769E-07 | 3.19314E-05 | 0.000415109 | -1.568879463 | 28.42403297 | RAB5C;TFRC;SRC;ARPC1A;CLTC;CBLB;ADRB2;WASL;RAB22A;EGFR;IGF1R;EEA1;CDC42;CYTH3;PARD6B;RAB11FIP1;KIF5B;EPS15;LDLR;SNX5;SMAD2;ARFGEF1;HSPA8;PRKCI;PDCD6IP;IST1;CAV2;CAV1;ARAP2;VPS37A;VPS37B;IGF2R;TGFBR1;TGFBR2;EHD1;RAB10;ZFYVE16;NEDD4;TRAF6;CAPZA2;CHMP4C;MDM2;MET;FGFR2;ARF6;SPG20 |
| MAPK signaling pathway_Homo sapiens_hsa04010 | 45/255 | 2.37278E-08 | 3.25598E-07 | 4.46397E-05 | 0.000501182 | -1.564972331 | 27.47562316 | ATF2;ZAK;NLK;STK4;DUSP16;EGFR;CRKL;ELK4;RAP1B;CDC42;PPP3CA;RPS6KA3;NRAS;MAPK8;MYC;MKNK2;MAPK1;PAK2;MAP3K2;MAP2K3;HSPA8;JUN;MAP3K1;CHUK;DUSP1;RRAS2;BRAF;FOS;GNG12;DUSP8;TGFBR1;DUSP7;CDC25B;TGFBR2;IL1A;TAOK1;TRAF6;RASA1;MAPKAPK2;RASA2;NF1;RAPGEF2;TAB2;CRK;FGFR2 |
| Chronic myeloid leukemia_Homo sapiens_hsa05220 | 21/73 | 2.93371E-08 | 3.81383E-07 | 1.72682E-05 | 0.000250896 | -1.348210794 | 23.38392388 | CDKN1A;CDKN1B;CHUK;BRAF;CBLB;PIK3CB;PIK3R1;TGFBR1;RUNX1;TGFBR2;CRKL;NRAS;CDK6;PIK3CA;CCND1;MYC;MDM2;E2F2;MAPK1;E2F3;CRK |
| TNF signaling pathway_Homo sapiens_hsa04668 | 26/110 | 6.08046E-08 | 7.50937E-07 | 3.42556E-05 | 0.000422984 | -1.403086524 | 23.31312479 | ATF2;TNFAIP3;CXCL1;NOD2;PIK3R1;PIK3CB;CXCL3;PTGS2;CXCL5;MAPK8;CREB3L2;MAPK1;JUNB;MAP2K3;EDN1;JUN;JAG1;CHUK;LIF;DAB2IP;CFLAR;FOS;CREB1;PIK3CA;TAB3;TAB2 |
| Bladder cancer_Homo sapiens_hsa05219 | 15/41 | 8.04703E-08 | 9.46484E-07 | 2.91535E-05 | 0.00040005 | -0.891028564 | 14.55528805 | CDKN1A;SRC;BRAF;THBS1;EGFR;VEGFA;NRAS;CCND1;CDH1;MYC;MDM2;E2F2;MAPK1;E2F3;HBEGF |
| Shigellosis_Homo sapiens_hsa05131 | 19/65 | 1.0021E-07 | 1.12508E-06 | 3.59622E-05 | 0.000422984 | -0.984198069 | 15.86133438 | ITGB1;CHUK;FBXW11;ROCK2;SRC;ARPC1A;NOD2;WASL;ACTB;CRKL;CDC42;MAPK8;CTTN;MAPK1;ITGA5;PFN1;CRK;VCL;CD44 |
| Rap1 signaling pathway_Homo sapiens_hsa04015 | 38/211 | 1.66939E-07 | 1.79278E-06 | 0.000121014 | 0.001153385 | -1.408555998 | 21.98141249 | ITGB1;SRC;CTNND1;GNAI3;LPAR2;PIK3R1;PIK3CB;EFNA5;MLLT4;THBS1;ADCY6;ACTB;EGFR;CRKL;IGF1R;RAP1B;CDC42;PARD6B;NRAS;CDH1;MAPK1;PLCG1;MAP2K3;PRKCI;INSR;BRAF;VEGFA;TIAM1;PIK3CA;ADORA2B;GNAS;RAPGEF2;CTNNB1;PFN1;CRK;MET;FGFR2;EPHA2 |
| Hepatitis B_Homo sapiens_hsa05161 | 30/146 | 1.74367E-07 | 1.79453E-06 | 8.45482E-05 | 0.000907974 | -1.350119213 | 21.01069258 | ATF2;CDKN1A;DDX3X;CDKN1B;YWHAB;SRC;PTEN;PIK3R1;PIK3CB;NRAS;MAPK8;CCND1;MYC;CREB3L2;E2F2;MAPK1;E2F3;JUN;MAP3K1;APAF1;CHUK;FOS;HSPG2;YWHAZ;TGFBR1;CREB1;CDK6;PIK3CA;CDK2;IFNAR1 |
| Apoptosis_Homo sapiens_hsa04210 | 29/140 | 2.32302E-07 | 2.29514E-06 | 9.78486E-05 | 0.001007025 | -1.344380199 | 20.5357156 | XIAP;PIK3R1;PIK3CB;ACTB;LMNB1;NRAS;MAPK8;BCL2L11;CTSL;CAPN2;PMAIP1;CASP2;MAPK1;CTSD;SPTAN1;CTSC;CTSB;MCL1;JUN;APAF1;CHUK;PDPK1;DAB2IP;TNFRSF10B;CFLAR;FOS;PTPN13;PIK3CA;ATM |
| AGE-RAGE signaling pathway in diabetic complications_Homo sapiens_hsa04933 | 23/101 | 7.04872E-07 | 6.44827E-06 | 0.000155094 | 0.00141882 | -1.31837144 | 18.67506123 | SMAD2;EDN1;JUN;CDKN1B;SERPINE1;PIK3R1;PIK3CB;F3;TGFBR1;TGFBR2;VEGFA;CDC42;IL1A;THBD;NRAS;MAPK8;CCND1;PIK3CA;COL4A2;COL4A1;PIM1;MAPK1;PLCG1 |
| Pancreatic cancer_Homo sapiens_hsa05212 | 18/66 | 6.80538E-07 | 6.44827E-06 | 0.000121409 | 0.001153385 | -1.087114109 | 15.43743639 | SMAD2;CHUK;TGFA;BRAF;PIK3CB;PIK3R1;TGFBR1;EGFR;TGFBR2;VEGFA;CDC42;MAPK8;CDK6;PIK3CA;CCND1;E2F2;MAPK1;E2F3 |
| Axon guidance_Homo sapiens_hsa04360 | 26/127 | 1.21888E-06 | 1.07522E-05 | 0.000260526 | 0.002115869 | -1.082876274 | 14.74615607 | ITGB1;GSK3B;NRP1;SEMA3C;ROCK2;GNAI3;EFNA5;CDC42;PPP3CA;NRAS;ABLIM1;CFL2;PAK6;PLXNA1;MAPK1;PAK2;EPHA4;ARHGEF12;SEMA4D;SEMA4B;LIMK2;SEMA4C;RASA1;EPHA1;MET;EPHA2 |
| Non-small cell lung cancer_Homo sapiens_hsa05223 | 16/56 | 1.4266E-06 | 1.21507E-05 | 0.000183281 | 0.001616797 | -0.911760645 | 12.27249335 | PDPK1;TGFA;BRAF;PIK3CB;PIK3R1;FOXO3;STK4;EGFR;NRAS;CDK6;PIK3CA;CCND1;E2F2;MAPK1;E2F3;PLCG1 |
| TGF-beta signaling pathway_Homo sapiens_hsa04350 | 20/84 | 1.75931E-06 | 1.4485E-05 | 0.000249208 | 0.002115869 | -0.995769726 | 13.19453293 | ACVR1;SMAD2;TGIF1;BMPR2;FST;CUL1;ACVR1B;THBS1;TGFBR1;ACVR2A;TGFBR2;SMAD7;PPP2CB;ZFYVE16;RPS6KB1;SP1;MYC;MAPK1;E2F5;BMPR1A |
| Transcriptional misregulation in cancer_Homo sapiens_hsa05202 | 32/180 | 2.05816E-06 | 1.63989E-05 | 0.000500675 | 0.003637259 | -1.043074923 | 13.65771029 | DDX5;CDKN1A;CCNT2;CDKN1B;KMT2A;DOT1L;JMJD1C;IGF1R;ELK4;HOXA9;CCND2;PLAU;MYC;NFKBIZ;WHSC1;JUP;H3F3B;FUS;IGFBP3;HMGA2;KLF3;PBX1;RUNX1;TGFBR2;GOLPH3;SPINT1;BCL6;SP1;BMP2K;MDM2;ATM;MET |
| Glioma_Homo sapiens_hsa05214 | 17/65 | 2.61056E-06 | 1.98705E-05 | 0.000283674 | 0.002189605 | -0.929807675 | 11.95355757 | CDKN1A;PTEN;TGFA;BRAF;PIK3CB;PIK3R1;EGFR;IGF1R;NRAS;CDK6;PIK3CA;CCND1;MDM2;E2F2;MAPK1;E2F3;PLCG1 |
| Endometrial cancer_Homo sapiens_hsa05213 | 15/52 | 2.65476E-06 | 1.98705E-05 | 0.000265554 | 0.002115869 | -0.632105033 | 8.115695849 | GSK3B;PDPK1;PTEN;BRAF;PIK3CB;PIK3R1;FOXO3;EGFR;NRAS;PIK3CA;CCND1;CDH1;MYC;CTNNB1;MAPK1 |
| Signaling pathways regulating pluripotency of stem cells_Homo sapiens_hsa04550 | 27/142 | 3.42522E-06 | 2.46384E-05 | 0.000553718 | 0.003907667 | -1.005649004 | 12.65543484 | GSK3B;BMPR2;RIF1;PIK3R1;PIK3CB;ACVR1B;IGF1R;NRAS;MYC;SMARCAD1;MAPK1;SKIL;ACVR1;SMAD2;ZFHX3;PCGF3;FZD6;LIF;WNT7A;KLF4;ACVR2A;REST;PIK3CA;CTNNB1;IL6ST;FGFR2;BMPR1A |

| | | | | | | | | |
|---|---|---|---|---|---|---|---|---|
| Thyroid hormone signaling pathway_Homo sapiens_hsa04919 | 24/118 | 3.49127E-06 | 2.46384E-05 | 0.000482611 | 0.003612273 | -0.949245146 | 11.9274983 | PFKFB2;NOTCH2;MED1;GSK3B;NOTCH1;PDPK1;SRC;SLC2A1;ATP1B3;PIK3R1;PIK3CB;HIF1A;ACTB;MED13L;NRAS;MED13;CCND1;PIK3CA;MYC;MDM2;CTNNB1;MAPK1;ITGAV;PLCG1 |
| HTLV-I infection_Homo sapiens_hsa05166 | 40/258 | 4.27833E-06 | 2.93541E-05 | 0.00119118 | 0.007355539 | -1.068700316 | 13.2112168 | ATF2;NRP1;GSK3B;CDKN1A;SLC2A1;XIAP;PIK3R1;PIK3CB;ADCY6;ETS2;ELK4;PPP3CA;NRAS;CCND3;ZFP36;MAPK8;CCND2;CCND1;MYC;CDC27;E2F2;E2F3;MYBL1;SMAD2;JUN;MAP3K1;CHUK;FZD6;RRAS2;WNT7A;FOS;TGFBR1;TGFBR2;DLG1;CREB1;PIK3CA;CANX;TBPL1;CTNNB1;ATM |
| Cell cycle_Homo sapiens_hsa04110 | 24/124 | 8.57441E-06 | 5.724E-05 | 0.000890643 | 0.005945647 | -0.823124878 | 9.603174245 | SMAD2;YWHAE;GSK3B;CDKN1A;CDKN1B;YWHAB;CUL1;YWHAZ;CDC25A;CDC25B;CCND3;WEE1;CCND2;CDK6;CCND1;MYC;CDC27;CDK2;MDM2;E2F2;E2F3;ATM;E2F5;YWHAG |
| Melanoma_Homo sapiens_hsa05218 | 17/71 | 9.54266E-06 | 6.20273E-05 | 0.000675097 | 0.004631913 | -0.698982624 | 8.080056076 | CDKN1A;PTEN;BRAF;PIK3CB;PIK3R1;EGFR;IGF1R;NRAS;CDK6;PIK3CA;CCND1;CDH1;MDM2;E2F2;MAPK1;E2F3;MET |
| Viral carcinogenesis_Homo sapiens_hsa05203 | 33/205 | 1.29978E-05 | 8.23195E-05 | 0.001831958 | 0.010283946 | -1.015692047 | 11.4272763 | YWHAE;ATF2;CDKN1A;CDKN1B;DDX3X;YWHAB;SRC;UBE3A;CHD4;PIK3R1;PIK3CB;CDC42;NRAS;CCND3;CCND2;CCND1;CREB3L2;PMAIP1;MAPK1;YWHAG;JUN;ACTN1;YWHAZ;DLG1;CREB1;CDK6;PIK3CA;MAPKAPK2;RASA2;CDK2;MDM2;TBPL1;IL6ST |
| Neurotrophin signaling pathway_Homo sapiens_hsa04722 | 23/120 | 1.54871E-05 | 9.56329E-05 | 0.001267687 | 0.007455209 | -0.849623447 | 9.410006902 | YWHAE;GSK3B;JUN;MAP3K1;PDPK1;GAB1;FRS2;BRAF;PIK3R1;PIK3CB;FOXO3;CRKL;RAP1B;CDC42;RPS6KA3;NRAS;MAPK8;PIK3CA;TRAF6;MAPKAPK2;MAPK1;PLCG1;CRK |
| mTOR signaling pathway_Homo sapiens_hsa04150 | 15/60 | 1.81443E-05 | 0.000109309 | 0.000947126 | 0.006156317 | -0.455271233 | 4.970264956 | CAB39;PDPK1;PTEN;BRAF;PIK3CB;PIK3R1;HIF1A;VEGFA;RPS6KA3;PIK3CA;RPS6KB1;DDIT4;MAPK1;ULK1;RICTOR |
| ECM-receptor interaction_Homo sapiens_hsa04512 | 18/82 | 1.90777E-05 | 0.000112195 | 0.001133166 | 0.007176717 | -0.466859551 | 5.073357179 | ITGB1;SDC4;ITGA3;LAMA3;LAMC2;LAMC1;HSPG2;THBS1;COL4A2;COL4A1;DAG1;SDC1;ITGA6;ITGAV;ITGA5;ITGB6;AGRN;CD44 |
| Tight junction_Homo sapiens_hsa04530 | 25/139 | 2.12046E-05 | 0.000121803 | 0.001737595 | 0.00998107 | -0.697128336 | 7.50200094 | ZAK;SRC;PTEN;GNAI3;PPP2R2A;F11R;CLDN1;MLLT4;ACTB;CDC42;PPP2CB;NRAS;PARD6B;SPTAN1;PRKCI;ACTN1;SHROOM2;RRAS2;ASH1L;TJP1;CLDN4;CTTN;MYH9;CTNNB1;AMOTL1 |
| Colorectal cancer_Homo sapiens_hsa05210 | 15/62 | 2.76556E-05 | 0.000155249 | 0.001257245 | 0.007455209 | -0.310903293 | 3.263141886 | SMAD2;GSK3B;JUN;BRAF;PIK3CB;FOS;PIK3R1;TGFBR1;TGFBR2;MAPK8;PIK3CA;CCND1;MYC;CTNNB1;MAPK1 |
| Ras signaling pathway_Homo sapiens_hsa04014 | 34/227 | 4.50103E-05 | 0.000241686 | 0.004412316 | 0.021796843 | -0.844630491 | 8.453584851 | RAB5C;PIK3R1;RASAL2;PIK3CB;EFNA5;STK4;MLLT4;EGFR;ETS2;IGF1R;RAP1B;CDC42;NRAS;MAPK8;PAK6;ABL2;MAPK1;PLCG1;PAK2;CHUK;INSR;GAB1;RRAS2;GNG12;VEGFA;TIAM1;PIK3CA;RASA1;RASA2;NF1;MET;FGFR2;EPHA2;ARF6 |
| HIF-1 signaling pathway_Homo sapiens_hsa04066 | 20/103 | 4.48285E-05 | 0.000241686 | 0.002248832 | 0.012343588 | -0.65963577 | 6.604713046 | EDN1;EGLN3;CDKN1A;CDKN1B;TFRC;INSR;SERPINE1;SLC2A1;PIK3R1;PIK3CB;HIF1A;EGFR;IGF1R;VEGFA;PIK3CA;RPS6KB1;MKNK2;HMOX1;MAPK1;PLCG1 |
| Estrogen signaling pathway_Homo sapiens_hsa04915 | 19/99 | 8.12813E-05 | 0.000427159 | 0.003209835 | 0.016868709 | -0.628222941 | 5.91634869 | ATF2;HSPA8;JUN;SRC;GNAI3;PIK3R1;PIK3CB;FOS;EGFR;ADCY6;HSP90B1;NRAS;CREB1;PIK3CA;SP1;CREB3L2;GNAS;MAPK1;HBEGF |
| Epithelial cell signaling in Helicobacter pylori infection_Homo sapiens_hsa05120 | 15/68 | 8.7276E-05 | 0.000449108 | 0.002739822 | 0.014711654 | -0.215159169 | 2.010971242 | JUN;CHUK;SRC;ADAM10;CXCL1;F11R;EGFR;CDC42;TJP1;ADAM17;MAPK8;ATP6V1B2;PLCG1;MET;HBEGF |
| Salmonella infection_Homo sapiens_hsa05132 | 17/86 | 0.000129813 | 0.000654364 | 0.00398761 | 0.02028925 | -0.295727655 | 2.646589389 | JUN;ROCK2;ARPC1A;CXCL1;FOS;WASL;CXCL3;KLC1;ACTB;CDC42;TJP1;IL1A;DYNC1LI2;MAPK8;MYH9;MAPK1;PFN1 |
| Acute myeloid leukemia_Homo sapiens_hsa05221 | 13/57 | 0.000178223 | 0.000880423 | 0.004024993 | 0.02028925 | -0.021571785 | 0.186217864 | JUP;CHUK;BRAF;PIK3CB;PIK3R1;RUNX1;NRAS;PIK3CA;CCND1;RPS6KB1;MYC;PIM1;MAPK1 |
| Lysosome_Homo sapiens_hsa04142 | 21/123 | 0.00020166 | 0.000976669 | 0.006620916 | 0.029733934 | -0.518521977 | 4.412064675 | CD164;ASAH1;IDUA;CLTC;M6PR;GNS;LITAF;IGF2R;LAPTM4B;LAPTM4A;LAMP1;NPC1;CTSL;AP1G1;LAMP2;PSAP;AP1S2;TPP1;CTSD;CTSC;CTSB |
| Arrhythmogenic right ventricular cardiomyopathy (ARVC)_Homo sapiens_hsa05412 | 15/74 | 0.000237755 | 0.001129336 | 0.005438866 | 0.02534717 | -0.058784627 | 0.490514791 | DSP;ITGB1;JUP;ITGA3;ACTN1;ACTB;GJA1;DAG1;CTNNB1;DSG2;ITGA6;ITGAV;ITGA5;ITGB6;DSC2 |
| Central carbon metabolism in cancer_Homo sapiens_hsa05230 | 14/67 | 0.000272531 | 0.001270099 | 0.005666994 | 0.025921249 | -0.096084779 | 0.788640488 | SLC2A1;PTEN;PIK3CB;PIK3R1;HIF1A;EGFR;GLS;SLC7A5;NRAS;PIK3CA;MYC;MAPK1;MET;FGFR2 |
| Lysine degradation_Homo sapiens_hsa00310 | 12/52 | 0.000278771 | 0.001275118 | 0.005189081 | 0.024648136 | 0.410892387 | -3.363203724 | KMT2D;KMT2A;NSD1;KMT2C;DOT1L;SETD7;PLOD2;ASH1L;WHSC1;COLGALT1;WHSC1L1;SUV420H1 |
| Toxoplasmosis_Homo sapiens_hsa05145 | 20/118 | 0.000312631 | 0.001378926 | 0.008498571 | 0.035578764 | -0.399308865 | 3.222617007 | MAP2K3;ITGB1;HSPA8;CHUK;PDPK1;LAMA3;GNAI3;XIAP;LAMC2;PIK3R1;PIK3CB;LAMC1;MAPK8;PIK3CA;TRAF6;PPIF;MAPK1;TAB2;ITGA6;LDLR |
| Leukocyte transendothelial migration_Homo sapiens_hsa04670 | 20/118 | 0.000312631 | 0.001378926 | 0.008498571 | 0.035578764 | -0.341718889 | 2.757837861 | ITGB1;ROCK2;ACTN1;CTNND1;GNAI3;MSN;PIK3R1;PIK3CB;F11R;CLDN1;MLLT4;ACTB;RAP1B;CDC42;CLDN4;PIK3CA;CTNNB1;PLCG1;EZR;VCL |
| Choline metabolism in cancer_Homo sapiens_hsa05231 | 18/101 | 0.000326078 | 0.001413004 | 0.007903554 | 0.034248736 | -0.356354411 | 2.860946693 | JUN;CHKA;SLC44A1;SLC44A2;PDPK1;PIK3R1;PIK3CB;FOS;WASL;HIF1A;EGFR;NRAS;MAPK8;PIK3CA;RPS6KB1;SP1;MAPK1;PLCG1 |
| Steroid biosynthesis_Homo sapiens_hsa00100 | 6/20 | 0.000351231 | 0.001495761 | 0.004999134 | 0.02421149 | 3.634906049 | -28.91228017 | SQLE;SOAT1;SC5D;DHCR24;MSMO1;HSD17B7;LSS |
| Chagas disease (American trypanosomiasis)_Homo sapiens_hsa05142 | 18/104 | 0.000471661 | 0.001974582 | 0.010181674 | 0.040562476 | -0.255905891 | 1.96004701 | SMAD2;JUN;CHUK;SERPINE1;GNAI3;PPP2R2A;PIK3R1;CFLAR;PIK3CB;FOS;TGFBR1;TGFBR2;PPP2CB;MAPK8;PIK3CA;TRAF6;GNAS;MAPK1 |
| Pathogenic Escherichia coli infection_Homo sapiens_hsa05130 | 12/55 | 0.00048398 | 0.001992384 | 0.007565169 | 0.033367799 | 0.617894868 | -4.716680114 | CDC42;ITGB1;CTTN;ROCK2;CDH1;ARPC1A;CTNNB1;WASL;EZR;CLDN1;YWHAZ;ACTB |
| Prolactin signaling pathway_Homo sapiens_hsa04917 | 14/72 | 0.000591906 | 0.002396733 | 0.009660795 | 0.039118301 | 0.055957736 | -0.415887037 | GSK3B;SRC;PIK3CB;FOS;PIK3R1;FOXO3;NRAS;MAPK8;CCND2;PIK3CA;CCND1;MAPK1;SOCS6;SOCS5 |
| Thyroid cancer_Homo sapiens_hsa05216 | 8/29 | 0.000821581 | 0.003273073 | 0.008921976 | 0.0367288 | 1.896541791 | -13.47356358 | NRAS;CCND1;CDH1;MYC;CCDC6;CTNNB1;MAPK1;BRAF |
| Insulin resistance_Homo sapiens_hsa04931 | 18/109 | 0.00083937 | 0.003290863 | 0.015114145 | 0.058331152 | -0.218042742 | 1.544366031 | MGEA5;GSK3B;PDPK1;INSR;GFPT1;SLC2A1;PTEN;PIK3R1;PIK3CB;PTPRF;PPP1CB;RPS6KA3;MAPK8;CREB1;PIK3CA;RPS6KB1;CREB3L2;OGT |
| Sphingolipid signaling pathway_Homo sapiens_hsa04071 | 19/120 | 0.001033158 | 0.003926001 | 0.018296747 | 0.067747903 | -0.132865011 | 0.913464867 | CERS3;ASAH1;ASAH2;CERS6;SGMS1;ROCK2;PDPK1;PTEN;GNAI3;PPP2R2A;PIK3R1;PIK3CB;PPP2CB;NRAS;SGPL1;MAPK8;PIK3CA;MAPK1;CTSD |
| Fc gamma R-mediated phagocytosis_Homo sapiens_hsa04666 | 16/93 | 0.00101804 | 0.003926001 | 0.015592226 | 0.059250459 | -0.07238376 | 0.498715135 | MYO10;LIMK2;ARPC1A;PIK3CB;PIK3R1;WASL;CRKL;CDC42;MARCKS;PIK3CA;RPS6KB1;CFL2;MAPK1;PLCG1;CRK;ARF6 |
| T cell receptor signaling pathway_Homo sapiens_hsa04660 | 17/104 | 0.001297847 | 0.004784598 | 0.01938008 | 0.070395291 | -0.055072504 | 0.366069628 | GSK3B;JUN;CHUK;PDPK1;CBLB;PIK3R1;PIK3CB;FOS;CDC42;PPP3CA;NRAS;DLG1;PIK3CA;PAK6;MAPK1;PLCG1;PAK2 |
| VEGF signaling pathway_Homo sapiens_hsa04370 | 12/61 | 0.001283157 | 0.004784598 | 0.014808902 | 0.058060298 | 0.480669237 | -3.200503427 | CDC42;PPP3CA;NRAS;PIK3CA;SRC;MAPKAPK2;MAPK1;PIK3CB;PIK3R1;PLCG1;PTGS2;VEGFA |
| Osteoclast differentiation_Homo sapiens_hsa04380 | 20/132 | 0.00135099 | 0.004907274 | 0.023024077 | 0.081414528 | -0.130051061 | 0.859236607 | JUN;CHUK;PIK3R1;PIK3CB;FOS;TGFBR1;FOSL2;TGFBR2;IL1A;CYLD;PPP3CA;MAPK8;CREB1;PIK3CA;TRAF6;MAPK1;TAB2;SQSTM1;JUNB;IFNAR1 |
| Hepatitis C_Homo sapiens_hsa05160 | 20/133 | 0.001483934 | 0.005312053 | 0.024534224 | 0.084166017 | -0.093154981 | 0.606723867 | GSK3B;CDKN1A;CHUK;CD81;PDPK1;BRAF;PPP2R2A;PIK3R1;PIK3CB;CLDN1;EGFR;PPP2CB;NRAS;CLDN4;MAPK8;PIK3CA;TRAF6;MAPK1;LDLR;IFNAR1 |
| Epstein-Barr virus infection_Homo sapiens_hsa05169 | 27/202 | 0.001558921 | 0.005423287 | 0.034247317 | 0.114311991 | -0.218988687 | 1.415490661 | YWHAE;ATF2;GSK3B;CDKN1A;CDKN1B;YWHAB;TNFAIP3;PIK3R1;PIK3CB;MAPK8;MYC;PLCG1;YWHAG;MAP2K3;HSPA8;JUN;CHUK;YWHAZ;PIK3CA;NEDD4;TRAF6;CDK2;MDM2;TBPL1;POLR3G;TAB2;CD44 |
| AMPK signaling pathway_Homo sapiens_hsa04152 | 19/124 | 0.001538745 | 0.005423287 | 0.02399309 | 0.083468917 | -0.026398908 | 0.170980134 | PFKFB2;CAB39;PDPK1;INSR;PPP2R2A;PIK3R1;PIK3CB;HMGCR;FOXO3;IGF1R;RAB10;PPP2CB;CREB1;CCND1;PIK3CA;RPS6KB1;RAB14;CREB3L2;ULK1 |
| Progesterone-mediated oocyte maturation_Homo sapiens_hsa04914 | 16/98 | 0.001800252 | 0.006175864 | 0.023072943 | 0.081414528 | 0.04619024 | -0.291914403 | GNAI3;BRAF;PIK3R1;PIK3CB;CDC25A;ADCY6;CDC25B;IGF1R;RPS6KA3;MAPK8;PIK3CA;CDC27;CDK2;MAPK1;CPEB2;CPEB4 |
| Measles_Homo sapiens_hsa05162 | 20/136 | 0.001951458 | 0.006602878 | 0.029520067 | 0.099882965 | -0.041893382 | 0.261380289 | HSPA8;GSK3B;CDKN1B;CHUK;TNFRSF10B;MSN;TNFAIP3;CBLB;PIK3R1;PIK3CB;IL1A;CCND3;CCND2;CDK6;CCND1;PIK3CA;TRAF6;CDK2;TAB2;IFNAR1 |
| Dorso-ventral axis formation_Homo sapiens_hsa04320 | 7/27 | 0.002567854 | 0.008571079 | 0.018376961 | 0.067747903 | 2.822802699 | -16.83712848 | NOTCH2;NOTCH1;MAPK1;CPEB2;EGFR;ETS2;CPEB4 |
| Wnt signaling pathway_Homo sapiens_hsa04310 | 20/142 | 0.003265917 | 0.010755754 | 0.041726875 | 0.133851145 | -0.000322469 | 0.00184588 | GSK3B;JUN;FBXW11;ROCK2;CSNK1A1;FZD6;CUL1;WNT7A;NLK;LRP6;VANGL1;PPP3CA;CCND3;MAPK8;CCND2;CCND1;DAAM1;TBL1XR1;MYC;CTNNB1 |
| Oocyte meiosis_Homo sapiens_hsa04114 | 18/123 | 0.003373823 | 0.010964923 | 0.038994164 | 0.126731032 | 0.103271055 | -0.58778878 | YWHAE;FBXW11;YWHAB;CUL1;YWHAZ;ADCY6;IGF1R;PPP1CB;PPP3CA;PPP2CB;RPS6KA3;SLK;CDC27;CDK2;MAPK1;CPEB2;YWHAG;CPEB4 |
| Chemokine signaling pathway_Homo sapiens_hsa04062 | 24/187 | 0.004659459 | 0.014946575 | 0.062124819 | 0.187132076 | 0.043148137 | -0.231656136 | GSK3B;CHUK;ROCK2;SRC;GNAI3;BRAF;CXCL1;PPBP;PIK3R1;PIK3CB;WASL;FOXO3;GNG12;CXCL3;ADCY6;CXCL5;CRKL;RAP1B;CDC42;TIAM1;NRAS;PIK3CA;MAPK1;CRK |
| Ubiquitin mediated proteolysis_Homo sapiens_hsa04120 | 19/137 | 0.004836647 | 0.015122174 | 0.051957067 | 0.160417443 | 0.209970001 | -1.119462125 | UBE2H;MAP3K1;FBXW7;FBXW11;UBA6;HUWE1;CUL1;SYVN1;XIAP;UBE3A;CBLB;UBE2G1;UBE2Z;UBE2J1;TRAF6;NEDD4;CDC27;MDM2;UBE2K |
| Inositol phosphate metabolism_Homo sapiens_hsa00562 | 12/71 | 0.004827325 | 0.015122174 | 0.037125343 | 0.122266131 | 0.737953492 | -3.935847496 | PIKFYVE;MTMR3;PIK3CA;SYNJ1;IMPAD1;IPMK;PTEN;PIP4K2B;PIK3CB;PLCG1;MTMR4;PIK3C2A |
| Insulin signaling pathway_Homo sapiens_hsa04910 | 19/139 | 0.005664591 | 0.017489424 | 0.057741371 | 0.176075538 | 0.17046095 | -0.881883242 | GSK3B;PRKCI;PDPK1;INSR;CBLB;BRAF;PIK3R1;PIK3CB;PTPRF;CRKL;PPP1CB;NRAS;MAPK8;PIK3CA;RPS6KB1;PRKAR1A;MKNK2;MAPK1;CRK |
| NOD-like receptor signaling pathway_Homo sapiens_hsa04621 | 10/57 | 0.007425736 | 0.02264391 | 0.045040862 | 0.14082396 | 1.370991519 | -6.721702045 | MAPK8;CHUK;TRAF6;TNFAIP3;TAB3;MAPK1;CXCL1;TAB2;NOD2;HSP90B1 |

| | | | | | | | |
|---|---|---|---|---|---|---|---|
| N-Glycan biosynthesis_Homo sapiens_hsa00510 | 9/49 | 0.008017263 | 0.024149561 | 0.044989965 | 0.14082396 | 1.982903092 | -9.569803956 | GANAB;B4GALT1;RPN2;MAN1A2;MAN2A1;ALG2;MOGS;ALG11;MGAT2 |
| Oxytocin signaling pathway_Homo sapiens_hsa04921 | 20/158 | 0.010684994 | 0.031797513 | 0.09141489 | 0.243656351 | 0.172097532 | -0.78113606 | JUN;CDKN1A;PPP1R12A;ROCK2;SRC;GNAI3;PIK3R1;PIK3CB;FOS;PTGS2;EGFR;ADCY6;ACTB;PPP1CB;PPP3CA;NRAS;CCND1;PIK3CA;GNAS;MAPK1 |
| Phosphatidylinositol signaling system_Homo sapiens_hsa04070 | 14/98 | 0.011153753 | 0.032797345 | 0.073040314 | 0.214773304 | 0.500781397 | -2.251502762 | MTMR3;IPMK;PTEN;PIK3R1;PIK3CB;MTMR4;PIK3C2A;PIKFYVE;SYNJ1;PIK3CA;IMPAD1;PIP4K2B;PLCG1;CDS2 |
| GnRH signaling pathway_Homo sapiens_hsa04912 | 13/91 | 0.01414366 | 0.041099813 | 0.082001867 | 0.222576495 | 0.532211169 | -2.266415295 | MAP3K2;MAP2K3;JUN;MAP3K1;SRC;EGFR;ADCY6;CDC42;NRAS;MAPK8;GNAS;MAPK1;HBEGF |
| Platelet activation_Homo sapiens_hsa04611 | 16/122 | 0.015337686 | 0.04405126 | 0.098888852 | 0.257111016 | 0.326723739 | -1.36486959 | ITGB1;PRKCI;ARHGEF12;PPP1R12A;ROCK2;SRC;GNAI3;PIK3R1;PIK3CB;ADCY6;ACTB;PPP1CB;RAP1B;PIK3CA;GNAS;MAPK1 |
| B cell receptor signaling pathway_Homo sapiens_hsa04662 | 11/73 | 0.015845859 | 0.04498767 | 0.080513993 | 0.222576495 | 0.898462064 | -3.723987838 | GSK3B;PPP3CA;JUN;NRAS;PIK3CA;CHUK;CD81;MAPK1;PIK3CB;FOS;PIK3R1 |
| Phagosome_Homo sapiens_hsa04145 | 19/154 | 0.016246236 | 0.045600232 | 0.115603057 | 0.294370672 | 0.381930088 | -1.573511481 | ITGB1;RAB5C;TFRC;M6PR;THBS1;ACTB;EEA1;SEC61A2;DYNC1LI2;PIKFYVE;LAMP1;CTSL;LAMP2;CANX;ATP6V1B2;OLR1;ITGAV;ITGA5;VAMP3 |
| Longevity regulating pathway - multiple species_Homo sapiens_hsa04213 | 10/64 | 0.016481814 | 0.045741665 | 0.078652417 | 0.221095035 | 1.321294289 | -5.424570618 | HSPA8;NRAS;PIK3CA;RPS6KB1;INSR;PIK3CB;PIK3R1;FOXO3;ADCY6;IGF1R |
| cAMP signaling pathway_Homo sapiens_hsa04024 | 23/199 | 0.018307779 | 0.049216248 | 0.143389634 | 0.350665738 | 0.381743898 | -1.527139448 | JUN;PPP1R12A;ROCK2;GNAI3;RRAS2;BRAF;ATP1B3;PIK3R1;ADRB2;FOS;PIK3CB;ATP2B1;MLLT4;ADCY6;RAP1B;PPP1CB;TIAM1;MAPK8;CREB1;PIK3CA;CREB3L2;GNAS;MAPK1 |
| cGMP-PKG signaling pathway_Homo sapiens_hsa04022 | 20/167 | 0.018775269 | 0.049216248 | 0.131604825 | 0.328347393 | 0.423302444 | -1.682718114 | MEF2A;ATF2;PPP1R12A;ROCK2;INSR;GNAI3;ATP1B3;PIK3R1;PIK3CB;ADRB2;ATP2B1;ADCY6;PPP1CB;PPP3CA;CREB1;PIK3CA;CREB3L2;PPIF;MAPK1;MEF2D |
| Longevity regulating pathway - mammal_Homo sapiens_hsa04211 | 13/94 | 0.018201484 | 0.049216248 | 0.097428494 | 0.256008915 | 0.664505737 | -2.662177548 | ATF2;INSR;PIK3CB;PIK3R1;FOXO3;ADCY6;IGF1R;NRAS;CREB1;PIK3CA;RPS6KB1;CREB3L2;ULK1 |
| Pertussis_Homo sapiens_hsa05133 | 11/75 | 0.019128582 | 0.049216248 | 0.091741055 | 0.243656351 | 1.320935809 | -5.226377134 | ITGB1;IL1A;JUN;MAPK8;CFL2;TRAF6;GNAI3;MAPK1;ITGA5;FOS;CXCL5 |
| Regulation of lipolysis in adipocytes_Homo sapiens_hsa04923 | 9/56 | 0.018794875 | 0.049216248 | 0.081828727 | 0.222576495 | 1.65820651 | -6.589996284 | PIK3CA;INSR;GNAS;GNAI3;PIK3CB;ADRB2;PIK3R1;PTGS2;ADCY6 |
| Sphingolipid metabolism_Homo sapiens_hsa00600 | 8/47 | 0.018870542 | 0.049216248 | 0.077736163 | 0.221095035 | 2.828823498 | -11.23086264 | CERS3;UGCG;ASAH1;SGPL1;CERS6;ASAH2;SGMS1;KDSR |
| Circadian rhythm_Homo sapiens_hsa04710 | 6/30 | 0.019098123 | 0.049216248 | 0.070020506 | 0.208374277 | 5.423005362 | -21.46515113 | CREB1;FBXW11;CUL1;FBXL3;CSNK1D;CLOCK |
